# Vegfc–Vegfr3-Dependent Lymphatic Sprouting Requires Apelin Signaling

**DOI:** 10.64898/2026.04.01.715778

**Authors:** Franziska Lehne, Jean Eberlein, Enno Bockelmann, Ege Eryılmaz, Julian Malchow, Lina W. Weiss, Christian S. M. Helker

## Abstract

While Vegfc–Vegfr3 signaling is the primary driver of lymphangiogenesis, the role of G protein-coupled receptors (GPCRs), the most successful class of druggable targets in the human genome, remains far less understood. A previous study has implicated Apelin signaling and its receptor Apelin receptor (Aplnr), a class A GPCR, in lymphatic development, yet the underlying cellular and molecular mechanisms remain unclear. Here, we show that Apelin signaling is indispensable for Vegfc–Vegfr3-dependent lymphatic sprouting and promotes lymphatic endothelial cell (LEC) migration without affecting LEC specification. Loss of Apelin signaling resulted in defective sprouting of ECs from the posterior cardinal vein (PCV) and subsequent failure of lymphatic vessel formation. Conversely, Apelin overexpression induces ectopic endothelial extensions from the PCV, an effect that is suppressed by reducing Vegfr3 signaling. Mechanistically, we show that Vegfc signaling through ERK activation regulates Aplnr expression. We propose that the specific upregulation of Aplnrb in LECs renders them migration-competent, establishing Apelin signaling as a critical and non-redundant regulator of lymphatic sprouting. Overall, our results reveal a tightly coordinated signaling axis between growth factor and GPCR pathways that governs lymphatic endothelial behavior.

## Introduction

In vertebrates, lymphatic vessel development is evolutionarily conserved and is predominantly driven by vascular endothelial growth factor C (Vegfc) signaling through its receptor, fms-related receptor tyrosine kinase 4 (Flt4), also known as Vegfr3 [1, 2]. Vegfc–Flt4 signaling induces the expression of *prospero homeobox 1a* (*prox1a*), a key transcription factor that promotes lymphatic fate by suppressing the blood vascular lineage program [3–6]. Downstream of the Flt4 receptor, the MAPK/ERK and PI3K-AKT pathway play distinct but complementary roles during lymphatic development. ERK signaling is primarily involved in early lymphatic specification and migration, whereas AKT signaling functions predominantly at later stages [7, 8]. Studying lymphatic development in model organisms such as mice presents challenges due to experimental limitations. In contrast, zebrafish provide a powerful system for investigating vascular development *in vivo*, owing to their optical transparency and genetic accessibility [9, 10]. In zebrafish, the development of the trunk lymphatic system begins at approximately 32 hours post fertilization (hpf) with the specification of lymphatic endothelial cell (LEC) precursors from endothelial cells (ECs) of the posterior cardinal vein (PCV) [3]. By 36 hpf, LEC precursors, as well as venous ECs, begin sprouting from the PCV, a process dependent on Vegfc–Flt4 signaling [7, 11, 12]. Venous ECs undergo anastomosis with arterial intersegmental vessels (aISVs) to form venous ISVs (vISVs), while the lymphatic precursors migrate dorsally along aISVs toward the horizontal myoseptum [13, 14]. These LEC precursors later form the entire trunk lymphatic vasculature, including the thoracic duct (TD) [10].

Although Vegfc–Flt4 signaling is well established as the primary driver of lymphangiogenesis, additional regulatory mechanisms remain to be fully elucidated. Recent studies in mice and human lymphatic cell cultures suggest a role for G protein-coupled receptor (GPCR) signaling in lymphangiogenesis, primarily by modulating Vegfc–Flt4 signaling [15–19]. GPCRs regulate key physiological processes and are targeted by more than 30 % of FDA-approved drugs [20]. Among these, the Apelin receptor (Aplnr), a class A GPCR, has been extensively studied in the context of angiogenesis but remains poorly characterized in lymphatic development. In zebrafish, Apelin (Apln) signaling has been shown to regulate the migration of tip cells during ISV formation [21, 22]. Furthermore, Apelin signaling has been identified as a potential modulator of lymphatic vessel formation [23], although the underlying mechanisms remain unclear. In this study, we show that Apelin signaling is required for Vegfc–Flt4-dependent lymphatic sprouting in zebrafish. While Vegfc–Flt4 signaling governs both lymphatic specification and migration, we find that Apelin signaling acts specifically on the migratory behavior of LECs, without affecting their initial specification. Importantly, because Apelin selectively modulates LEC migration without interfering with Vegfc-driven specification, it emerges as a promising and more targeted therapeutic entry point for modulating lymphatic vessel growth in pathological contexts.

## Results

### Apelin-Aplnrb signaling is required for lymphatic development

In the developing blood vasculature, expression of the ligand Apelin is enriched within leading tip cells while the receptor is expressed in newly growing blood vessels but is absent from quiescent blood vessels [22, 24]. Here, we asked whether Apelin signaling might also affect venous and lymphatic sprouting. To test this, we imaged the vasculature in the trunk of *Tg(kdrl:EGFP)* larvae at 120 hpf (Figure 1A). We observed a reduced number of vISVs emerging from the PCV in *apln* heterozygous and *apln* homozygous mutant larvae (WT: 44.2 % ± 7.9 %, *apln +/-*: 34.2 % ± 11.1 %, *apln -/-*: 6.9 % ± 4.4 %, Figure 1A, C), indicating a defect in venous sprouting. This prompted us to investigate whether lymphatic sprouting is also affected in *apln* mutants. Therefore, we genetically labeled vascular ECs including lymphatic ECs using *Tg(fli1a:EGFP)* and assessed the presence of the TD. In wild type larvae, the TD is located between the dorsal aorta (DA) and the PCV at 120 hpf (Figure 1B, red arrowheads). As in mammals, there are two endogenous ligands for the Aplnr in the zebrafish genome, Apela [25, 26] and Apelin [27]. *apela* mutant larvae exhibited a normal TD formation (Figure 1B, D). In contrast, 80 % of *apln* mutants lack the TD and 13.3 % exhibit a partial TD (Figure 1B, D). Interestingly, *apln* heterozygous mutant larvae also showed defects in TD formation although to a lesser extent than homozygous mutants, suggesting a dose-dependent role for Apelin in TD formation (Figure S1A, B).

**Figure 1.**
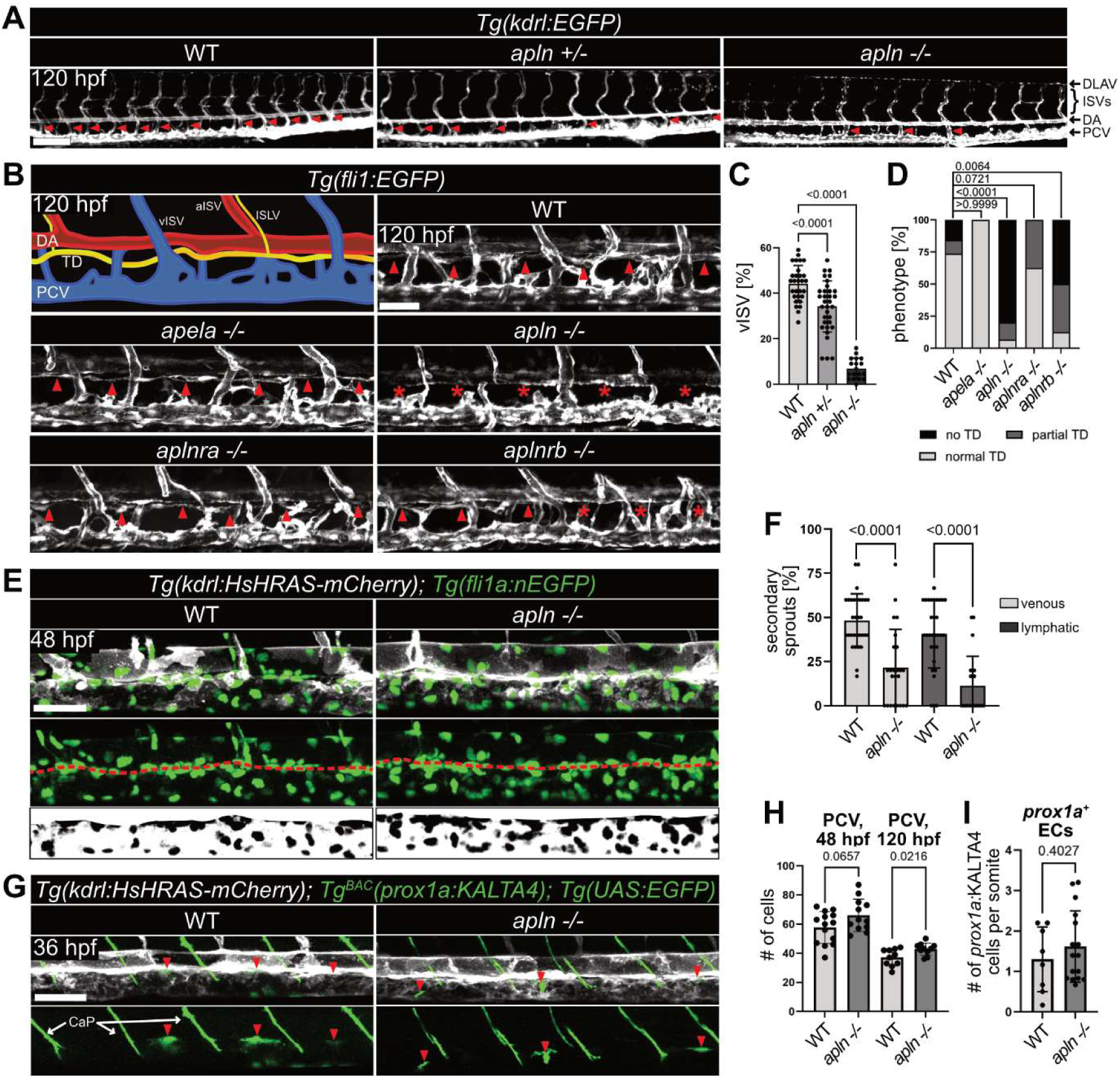
Apelin signaling is required for lymphatic development. **A** Fluorescence images of *Tg(kdrl:EGFP)* larvae at 120 hpf. *apln* mutant larvae exhibit fewer vISVs (red arrowheads). Quantification in C. **B** Schematic representation of the trunk vasculature at 120 hpf and confocal images of *Tg(fli1:EGFP)* larvae at 120 hpf. The location of the TD is indicated by red arrowheads. *apela* mutants form a TD (red arrowheads), while *apln* mutants do not form a TD (red asterisks). *aplnra* mutants form a TD (red arrowheads), while *aplnrb* mutants exhibit defects in TD formation (red asterisks). **C** Percentage of vISV formed across 15 somites corresponding to images in panel A. n(WT)= 29, n(*apln +/-*)= 33, n(*apln -/-*)= 18. *p*-values were calculated by One-way ANOVA with multiple comparison. **D** Quantification of the TD phenotype as fraction of total corresponding to images in B. TD phenotype is categorized as normal (light gray), partial (dark gray) or no TD (black). *p*-values were calculated by Fisher’s exact test. n(WT)= 19, n(*apela -/-*)= 2, n(*apln -/-*)= 15, n(*aplnra -/-*)= 16, n(*aplnrb -/-*)=8. **E** Confocal images of *Tg(kdrl:HsHRAS-mCherry); Tg(fli1a:nEGFP)* embryos at 48 hpf. Red dashed line indicates border between DA and PCV. Lower panel shows EC nuclei (black) in the PCV. **F** Percentage of venous (light gray) and lymphatic (dark gray) sprouts formed across five somites corresponding to images in panel E. *apln* mutants show significant reduction in both lymphatic and venous sprouting. n(WT)= 31, n(*apln -/-*) = 22. **G** Confocal images of the LEC marker *Tg^BAC^(prox1a:KALTA4); Tg(UAS:EGFP)* at 36 hpf. Red arrowheads indicate EGFP-positive cells in the PCV. Note, *prox1a:*KALTA4 is also expressed in the caudal primary motor neuron (CaP). **H** Quantification of the number of ECs in the PCV at 48 hpf (see panel E) and 120 hpf. Compared to their WT siblings, *apln* mutants exhibit more ECs in the PCV. **I** Quantification of *prox1a*:KALTA4-expressing cells (LECs) in the PCV per somite at 36 hpf corresponding to images in panel G. There is no significant difference in *apln* mutants compared to WT embryos. n(WT)= 8, n(*apln -/-*)= 16. Data are represented as mean ± SEM. Scale bars A: 150 µm, B, E and G: 50 µm. F,H,I: *p*-values were calculated by Student’s t-tests. hpf: hours post fertilization, DLAV: dorsal longitudinal anastomotic vessel, aISV/vISV: arterial/venous intersegmental vessel, DA: dorsal aorta, PCV: posterior cardinal vein, ISLV: intersegmental lymphatic vessel, TD: thoracic duct. See also related Figures S1 and S2.

The zebrafish genome contains two Apelin receptors: *aplnra* and *aplnrb* [28]. While both are differentially expressed, only loss of *aplnrb*, but not *aplnra*, partially mimicked the *apln* mutant phenotype (*aplnra -/-:* 37.5 % partial TD; *aplnrb -/-:* 37.5 % partial TD, 50 % no TD, Figure 1B, D). Given the presence of two paralogous receptors, we asked whether functional redundancy might account for this difference. Indeed, *aplnra -/- ; aplnrb -/-* double mutants completely lacked lymphatic sprouts and phenocopied the *apln* mutant phenotype (Figure S1C). Because Apelin receptor signaling through Apela has been shown to regulate early heart field formation, and receptor mutants display cardiovascular defects [28, 29], we focused our subsequent functional analyses primarily on *apln* mutants to avoid potential secondary effects arising from early cardiac abnormalities. Notably, while *apln* and *aplnra* mutants did not exhibit obvious gross morphological abnormalities, *apela*, *aplnrb*, and *aplnra/aplnrb* mutant embryos frequently developed cardiac edema, consistent with previously described roles of Apelin signaling during cardiac development [28, 29].

### *apln* mutants have an increased cell number in the PCV

Secondary sprouting starts at around 36 hpf, and by 48 hpf, venous sprouts have connected to arterial ISVs, while lymphatic ECs migrate to the horizontal myoseptum [13]. In *apln* mutant larvae, fewer sprouts emerged from the PCV, leading to a failure of vISV and TD formation (Figure 1A, B, S1A). To directly assess sprouting dynamics, we performed time-lapse imaging of *Tg(kdrl:HsHRAS-mCherry); Tg(fli1a:nEGFP)* embryos from 30 to 46 hpf (Video S1). While wild type siblings showed active EC sprouting from the PCV, *apln* mutant embryos failed to initiate sprouting (Video S1). To quantify this phenotype, we imaged wild type and *apln* mutant *Tg(kdrl:HsHRAS-mCherry); Tg(fli:nEGFP)* embryos at 48 hpf. Compared to wild type siblings, *apln* mutants exhibited significantly fewer venous (WT: 48.3 % ± 14.9 %, *apln -/-:* 21.6 % ± 22.3 %) and lymphatic sprouts (WT: 40.6 % ± 18.9 %, *apln -/-:* 11.4 % ± 16.2 %) that emerged from the PCV (Figure 1F), indicating a defect in secondary sprouting. Further analysis revealed an increased number of ECs in the PCV of *apln* mutants (WT: 57.6 ± 10.5, *apln -/-*: 66.0 ± 10.6, Figure 1E, H), while the diameter of the PCV remained unchanged (WT: 31.9 µm ± 3.4 µm, *apln -/-*: 31.2 µm ± 3.0 µm, Figure S1E). These effects were still visible at 120 hpf (Figure 1H, S1E). However, the number of ECs in the DA was not affected (Figure S1D), suggesting that the increase in EC number is specific to the PCV. Somite width increased over time and was unaffected in *apln* mutants indicating no general growth or developmental defect (Figure S1F). To exclude potentially higher mitotic activity in *apln* mutants as a reason for the higher cell number, we imaged wild type siblings and *apln* mutants expressing the cell cycle marker *Tg(kdrl:mVenus-gmnn)* [30] and counted *kdrl*:mVenus-gmnn-positive cells in the PCV at 36 hpf (Figure S2). Although this provided only a snapshot of proliferation at the time of imaging, we could not observe any changes in the number of mVenus-gmnn expressing cells that would explain the increased cell number in the PCV of *apln* mutant embryos.

In summary, *apln* mutants exhibit significantly reduced venous and lymphatic sprouting at 48 hpf and fail to form lymphatic vessels at 120 hpf. This sprouting defect is accompanied by an increased number of endothelial cells in the PCV, likely reflecting a failure of cells destined for secondary sprouting to migrate out at the correct developmental time.

### Lymphatic differentiation is not affected in *apln* mutants

Since *apln* mutants fail to form lymphatic vessels, we wondered whether LECs are specified in the PCV. To test this, we examined the expression of the LEC marker *Tg^BAC^(prox1a:KALTA4,4xUAS-E1B:TagRFP)* [31] (further referred to as *Tg^BAC^(prox1a:KALTA4)*) in *apln* mutant embryos (Figure 1G, I). Imaging of *apln* mutant embryos revealed normal *prox1a*:KALTA4 expression in the PCV at 36 hpf, indicating that LEC specification is not affected. These results suggest that Apelin signaling primarily regulates LEC migration rather than their differentiation. Notably, because the lymphatic phenotype is more severe than the venous phenotype, LECs appear to be particularly sensitive to the loss of Apelin signaling during this developmental window.

### Fibroblasts and ECs of the ISVs express *apln*

Previous studies have shown that Apelin-Aplnrb signaling regulates aISV migration during trunk angiogenesis [22]. During aISV development, Apelin is produced by neural precursors and the vasculature itself [21]. Here, we used the *Tg^BAC^(apln:Venus-PEST)* reporter line to examine the expression pattern during venous and lymphatic development at 48 hpf. As previously shown, *apln:*Venus-PEST was highly expressed in the dorsal ISVs and in neural progenitors [21] (Figure 2A, yellow arrowheads), while weak *apln*:Venus-PEST expression was observed in cells at the horizontal myoseptum (red arrowheads). Previous work has shown that these cells are fibroblasts and are expressing Vegfc to attract lymphatic cells to migrate along the midline [31, 32]. To test whether fibroblasts expressing *apln*:Venus-PEST also express *vegfc*, we analyzed *Tg^2BAC^(vegfc:Gal4FF); Tg(UAS:EGFP); Tg^BAC^(apln:Venus-PEST)* embryos at 36 hpf. We observed a partial overlap between *apln*:Venus-PEST and *vegfc*:Gal4FF expression (Figure 2B, red arrowheads), suggesting that some cells express both ligands and that the *apln*:Venus-PEST-expressing cells at the midline likely represent a subpopulation of fibroblasts.

**Figure 2.**
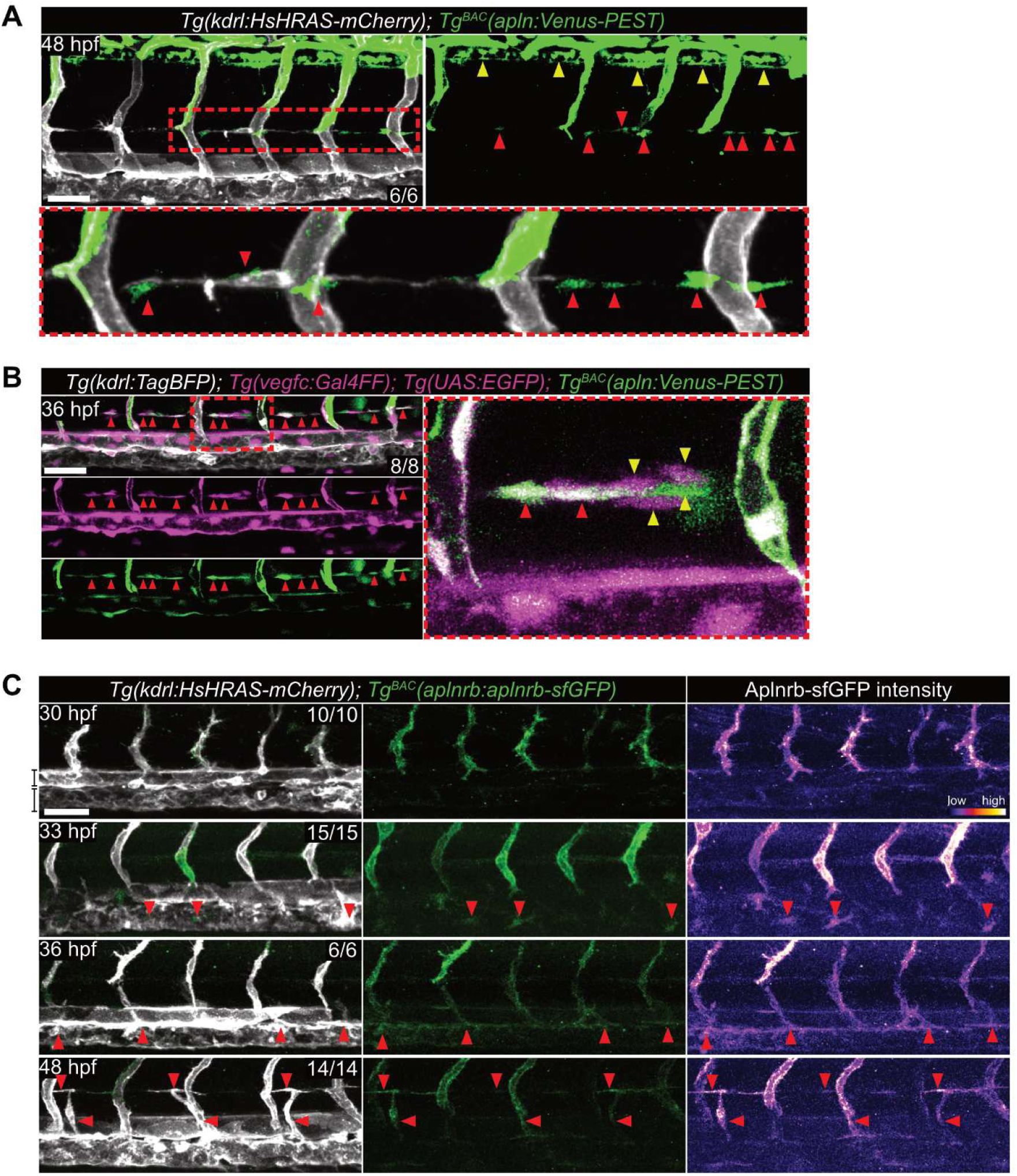
Fibroblasts and ECs of the ISVs express *apln* and sprouting LECs upregulate *aplnrb*. **A** Confocal image of a *Tg^BAC^(apln:Venus-PEST)* embryo at 48 hpf. *apln* is expressed in the neuronal tube (yellow arrowheads) and the dorsal part of arterial and venous ISVs. Magnification of boxed area is shown in the lower panel. Some *apln*-expressing cells locate at the horizontal myoseptum (red arrowheads). *kdrl:*HsHRAS-mCherry- positive cells (gray) migrate along the *apln*-expressing cells. **B** Confocal image of a *Tg(vegfc:Gal4FF); Tg(UAS:EGFP)*; *Tg^BAC^(apln:Venus-PEST)* embryo at 36 hpf. *vegfc* expression (magenta) can be seen in the DA and in a subpopulation of fibroblasts at the horizontal myoseptum. Magnification of boxed area from top panel is on the right. Cells at the horizontal myoseptum co-expressing *vegfc:*Gal4FF and *apln:*Venus-PEST are indicated by red arrowheads while cells expressing only one of the markers are indicated by yellow arrowheads. **C** Confocal images of *Tg(kdrl:HsHRAS-mCherry)*; *Tg^BAC^(aplnrb:aplnrb-TagRFP-sfGFP)* embryos at 30, 33, 36 and 48 hpf. Aplnrb-sfGFP fluorescence intensity is visualized with a fire LUT in the right panel. At 30 hpf, ISVs exhibit high expression of *aplnrb:*Aplnrb-sfGFP. At 33 hpf, single cells in the PCV show increased *aplnrb:*Aplnrb-sfGFP expression (red arrowheads). At 36 and 48 hpf, lymphatic sprouts and lymphatic precursors at the midline show high *aplnrb:*Aplnrb-sfGFP expression (red arrowheads). Scale bars: 50 µm. See also related Figure S3.

### Sprouting LECs upregulate *aplnrb* in a time- and cell type-specific manner

To investigate the Aplnrb protein expression in the PCV during secondary sprouting, we imaged *Tg^BAC^(aplnrb:aplnrb-TagRFP-sfGFP*) [22] (further referred to as *Tg^BAC^(aplnrb:aplnrb-sfGFP)*) embryos at different stages (Figure 2C). At 30 hpf, prior to LEC fate commitment, *aplnrb*:Aplnrb-GFP expression can be seen in the aISVs while no ECs in the DA or PCV show fluorescence. However, at 33 hpf, coinciding with the onset of LEC specification, individual cells within the PCV upregulate *aplnrb*:Aplnrb-GFP expression (Figure 2C, red arrowheads) displaying a spatial pattern reminiscent of previously described lymphatic angioblasts [33]. From 36 hpf onwards, robust *aplnrb*:Aplnrb-GFP expression was consistently detected in all secondary sprouts from the PCV (red arrowheads). To better understand the temporal and cellular context in which Apelin signaling may function during lymphatic development, we analyzed publicly available single-cell RNA sequencing (scRNA-seq) datasets from zebrafish [4, 34] (Figure S3). Within the EC cluster of the vascular atlas [34], *aplnrb* gene expression positively correlates with the pro-migratory gene *egfl7* [36], and several well-established pro-lymphatic genes, including the transcription factor *sox18* [37] and the receptor tyrosine kinases *tie1* [38] and *flt4* [1] (Figure S3A). In the lymphangiogenesis-specific scRNA-seq dataset [4], *aplnrb* expression peaks at 40 hpf and 3 dpf, then declines at later stages (4 and 5 dpf) (Figure S3B). This temporal pattern suggests that Aplnrb signaling is most relevant during early LEC specification and sprouting, rather than vessel maturation. In contrast, *flt4* remains robustly expressed throughout all stages of lymphatic development (Figure S3C). Cluster-level analysis further reveals that *aplnrb* is predominantly expressed in the pre-LEC cluster, which consists of early specified lymphatic endothelial cells that are not yet fully distinct from venous endothelial cells (VECs) (Figure S3D). By comparison, *flt4* expression is strongly expressed in differentiated LECs (Figure S3E). These data support a model in which Apelin signaling acts early and transiently to regulate progenitor migration, while Vegfc–Flt4 signaling plays a continuous role in LEC maturation and maintenance.

### Overexpression of Apelin induces ectopic endothelial extensions from the PCV

Given that *aplnrb* expression coincided with the timing of LEC sprouting from the PCV, we next investigated the effect of Aplnrb overactivation. Therefore, we ubiquitously overexpressed its ligand Apelin using a heat shock-inducible transgene (*Tg(hsp70l:apln)*), thereby saturating the tissue with uniformly distributed Apelin. To identify a potential developmental time window during which ECs are responsive to Apelin signaling, we induced Apelin overexpression at different developmental time points, (Figure 3A, B). We found that the overexpression of Apelin at 24 hpf resulted in the most pronounced and severe phenotype, suggesting that LECs are particularly sensitive to Apelin during the initial stages of secondary sprouting (Figure 3C-F). Similar to *apln* mutants, Apelin overexpression also resulted in an increased EC number in the PCV at 48 hpf (Figure S4A, B). Notably, Apelin overexpression also severely impacted venous sprouting (Figure 3D), supporting the idea that both lymphatic and venous ECs require precise regulation of Apelin levels and spatial distribution to ensure proper migratory behavior. High resolution confocal imaging further highlights the pronounced venous hypersprouting observed upon global Apelin overexpression (Figure 3F).

**Figure 3.**
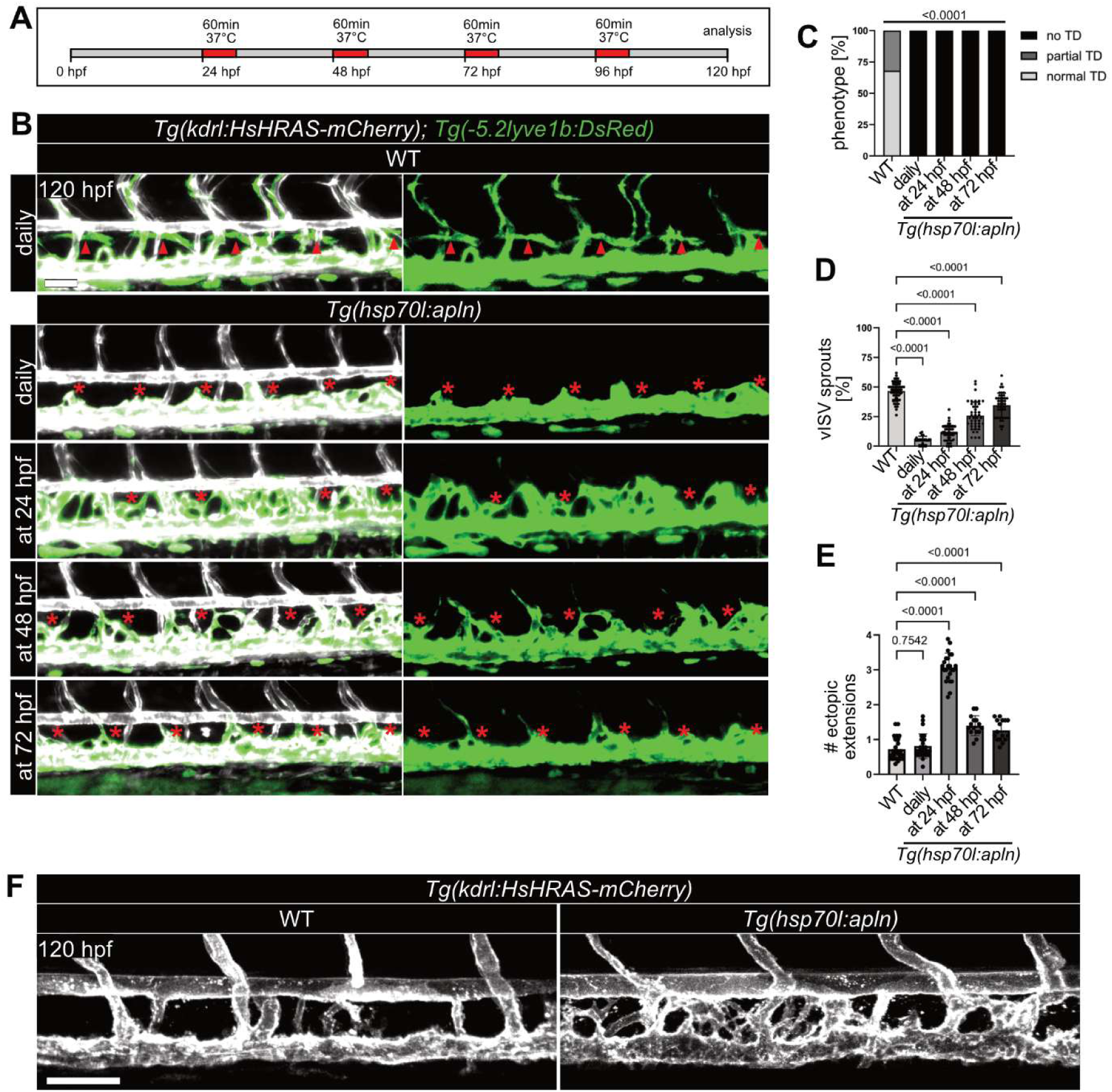
Apelin signaling induces ectopic endothelial extensions. **A** Schematic representation of the used heat shock protocol to temporally control induction of ubiquitous Apelin overexpression. Embryos were heat shocked either daily or once at 24 hpf, 48 hpf or 72 hpf at 37 °C. **B** Fluorescence images of *Tg(kdrl:EGFP)*; *Tg(-5.2lyve1b:DsRed); Tg(hsp70l:apln)* larvae at 120 hpf. Larvae with induced, ubiquitous *apln* overexpression completely lack lymphatic structures (red asterisks) independent of time of induction. **C** Quantification of TD phenotype. *p*-value was calculated by Fisher’s exact test. **D** Percentage of vISVs formed across 5 somites. vISV formation is reduced dependent on the time of induction of Apelin overexpression. **E** Number of ectopic endothelial extensions across six somites. Larvae overexpressing Apelin form more ectopic extensions from the PCV dependent on the time of induction. **F** High resolution confocal images of *Tg(kdrl:HsHRAS-mCherry); Tg(hsp70l:apln)* larvae at 120 hpf. Larvae were heat shocked at 24 hpf. Ubiquitous overexpression of Apelin leads to formation of ectopic endothelial extensions from the PCV. D and E: n(WT)= 25, n(daily)= 24, n(at 24 hpf)= 23, n(at 48 hpf)= 15, n(at 72 hpf)= 16. E: n(WT)= 82, n(daily)= 24, n(at 24 hpf)= 54, n(at 48 hpf)= 43, n(at 72 hpf)= 45. Scale bars: 50 µm. D and E: *p*-values were calculated by One-way ANOVA. See also related Figure S4.

### Vascular-derived Apelin can partially rescue TD formation in *apln* mutants

During the developmental window of lymphangiogenesis, we identified three potential sources of Apelin: fibroblasts, neural progenitors and ECs of the ISVs (Figure 2A, B). To determine which source of Apelin contributes functionally to lymphatic development, we performed rescue experiments by cell type-specific re-expression of Apelin in *apln* mutants (Figure 4A, B). First, we tested ubiquitous overexpression of Apelin using *Tg(hsp70l:apln)*. However, global overexpression of Apelin does not rescue TD formation in *apln* mutants (Figure 4B, C). Next, we tested if *vegfc*-expressing cells can rescue the *apln* mutant phenotype. Surprisingly, even though *vegfc:*Gal4 is expressed in arteries and in fibroblasts at the horizontal myoseptum, no rescue of TD formation was observed (Figure 4B, C). Furthermore, we tested whether neural progenitor-derived (*Tg(zic2b:Gal4)*) Apelin rescues secondary sprouting defects, as previously shown for arterial sprouting [21]. However, only 22,2 % larvae with neural progenitor-derived Apelin exhibited a TD at 120 hpf (Figure 4B, C). Lastly, we assessed whether Apelin from the vasculature (*Tg(fli1a:Gal4)*) could compensate for the loss of endogenous *apln.* Consistent with previous work showing that *apln* is transiently expressed in arterial ISVs during zebrafish vascular development [22], 58,3 % of larvae with vascular-derived Apelin showed a rescue of TD formation at 120 hpf (Figure 4B, C).

**Figure 4.**
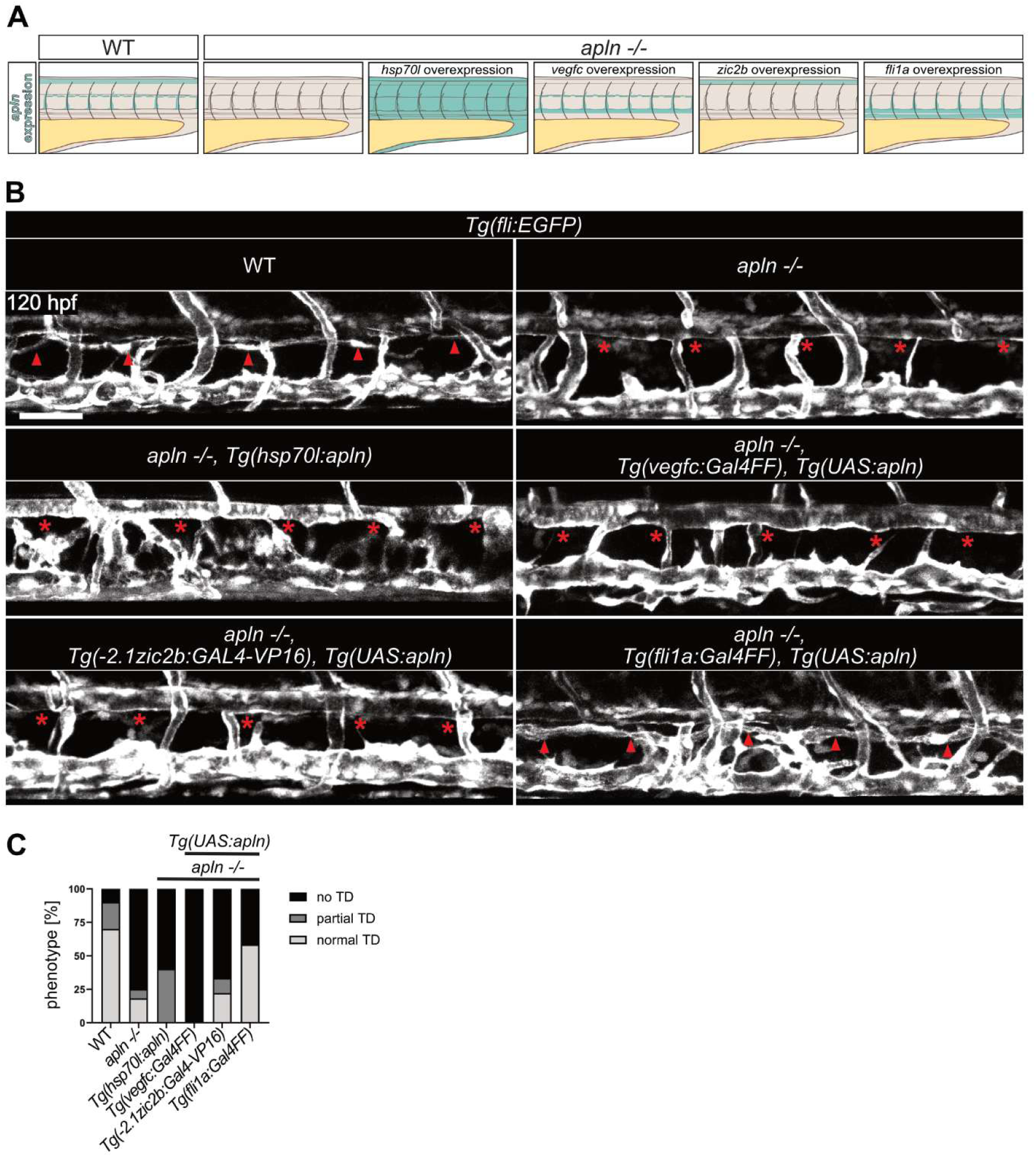
Vascular-derived Apelin partially rescues TD formation in *apln* mutants. **A** Schematic overview of the tissue-specific Apelin expression rescue experiments. **B** Confocal images of larvae expressing *Tg(fli:EGFP)* and the respective transgenes for rescue at 120 hpf. Siblings exhibit a TD (red arrowheads) while *apln* mutants fail to develop a TD (red asterisks). Larvae with *Tg(hsp70l:apln)* were heat shocked at 24 hpf leading to ubiquitous overexpression of Apelin, which is unable to rescue the lack of TD formation (red asterisks). *apln* expression in *vegfc*-expressing cells or neural progenitors fail to restore TD formation. Vascular expression of *apln* can restore TD formation in 7 out of 12 imaged larvae. **C** Quantification of TD formation of re-expressing *apln* in different tissues to rescue the *apln* mutant phenotype. n(WT)=50, n(*apln -/-*)= 44, rescues: n(*Tg(hsp70l:apln)*)= 5, n(*Tg(fli:Gal4FF)*)= 12, n*(Tg(- 2.1zic2b:Gal4-VP16)*):= 9, n(*Tg(vegfc:Gal4FF)*)= 4. Scale bars: 50 µm.

### Genetic interaction between Apelin and Vegfc signaling

Since the lymphatic phenotype of the *apln* mutants resembled those of loss of function of Vegfc signaling, we investigated whether Apelin functions upstream of, or in coordination with, the Vegfc–Flt4 signaling pathway. To address this, we performed whole-mount *in situ* hybridization (WISH) in *apln* mutant embryos at 32 hpf to assess the expression of *vegfc*, *flt4* and *etsrp*, a transcription factor that has been shown to regulate *flt4* expression during lymphangiogenesis [39]. The expression patterns in *apln* mutants were indistinguishable from wild type siblings (Figure 5A, S5), suggesting that Apelin signaling does not directly regulate Vegfc–Flt4 signaling at the transcriptional level. To further examine the functional interaction between Apelin and Vegfc–Flt4 signaling, we analyzed TD and vISV formation in *apln* and *flt4* double heterozygous mutant larvae at 120 hpf (Figure 5B, C, D). While single heterozygous mutants exhibited mild defects, the double heterozygous mutants phenocopied homozygous *apln* mutants and displayed more severe defects, including significant reduction in vISV formation and complete absence of the TD (Figure 5C, D). Together, these findings suggest that Apelin does not act upstream of Vegfc–Flt4 signaling but Apelin and Vegfc–Flt4 signaling genetically interact.

**Figure 5.**
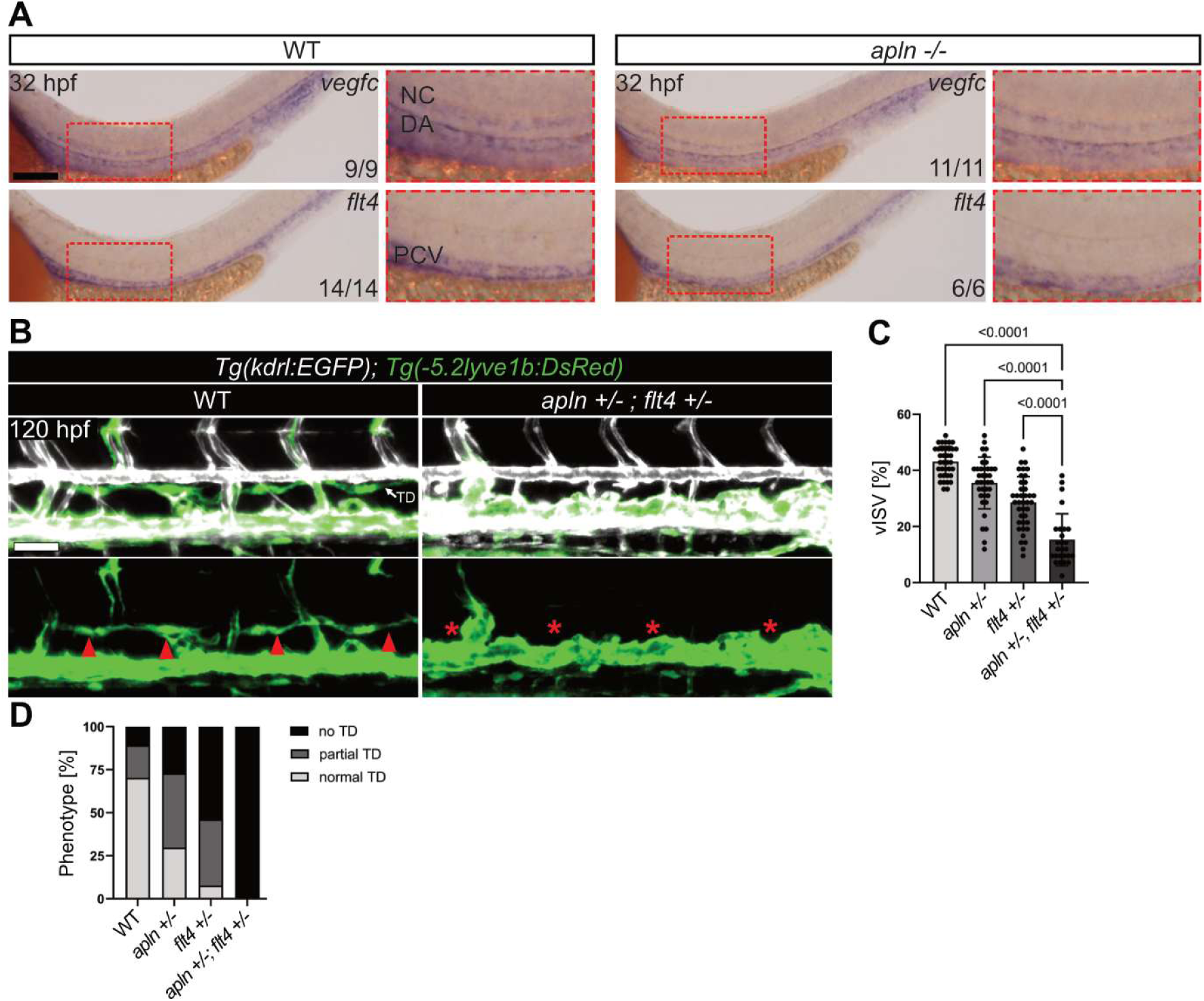
Genetic interaction between Apelin and Vegfc signaling. **A** Whole mount *in situ* hybridization against *vegfc* and *flt4* of *apln* mutants and siblings at 32 hpf. Compared to siblings, *apln* mutants exhibit a similar *vegfc* and *flt4* expression patterns. **B** Fluorescence images of *Tg(kdrl:EGFP); Tg(-5.2lyve1b:DsRed)* larvae at 120 hpf. *apln* and *flt4* double-heterozygous larvae do not form a TD (asterisks) and exhibit reduced vISV numbers. Quantifications in **C** and **D**. n(WT)= 36, n(*apln +/-*)= 37, n(*flt4 +/-*)= 39, n(*apln +/-, flt4 +/-*) = 26. D: *p*-values were calculated by One-way ANOVA with Multiple Comparison. NC: notochord, DA: dorsal aorta, PCV: posterior cardinal vein. Scale bars A: 100 µm, B: 50 µm.

### Overexpression of Vegfc induces *aplnrb* expression via ERK signaling

To further explore the interaction between Vegfc–Flt4 and Apelin signaling, we examined whether Vegfc overexpression modulates *aplnrb* expression. Using time-lapse imaging of *Tg^BAC^(prox1a:KALTA4); Tg(UAS:vegfc); Tg^BAC^(aplnrb:Venus-PEST)* embryos from 30 hpf, we observed a strong induction of *aplnrb:*Venus-PEST expression in the PCV (Figure 6A, B, Video S2). Interestingly, this upregulation of *aplnrb:*Venus-PEST expression was spatially restricted to the PCV, as *aplnrb*:Venus-PEST expression in the ISVs was slightly reduced in response to increased Vegfc signaling (Figure 6C), suggesting that Vegfc–Flt4 signaling regulates *aplnrb* expression specifically within the PCV. To confirm whether this effect is driven by physiological relevant Vegfc-expressing cells, we imaged *Tg(vegfc:Gal4FF); Tg(UAS:vegfc); Tg^BAC^(aplnrb:Venus-PEST)* embryos at 82 hpf. Overexpression of Vegfc in endogenously Vegfc-expressing cells was sufficient to upregulate *aplnrb:*Venus-PEST expression specifically in the PCV (Figure S6). To test whether Flt4 signaling is required for *aplnrb* upregulation during secondary sprouting, we next analyzed *Tg^BAC^(aplnrb:aplnrb-TagRFP-sfGFP)* expression in *flt4* mutant embryos. In wild type siblings, Aplnrb-sfGFP-positive cells appeared in the PCV from 33 hpf onwards and showed robust expression at 36 and 48 hpf, consistent with the timing of secondary sprouting from the PCV (Figure 6D, E). In contrast, *flt4* mutant embryos showed a strong reduction of Aplnrb-sfGFP-positive cells in the PCV at these stages (Figure 6D, E).

**Figure 6.**
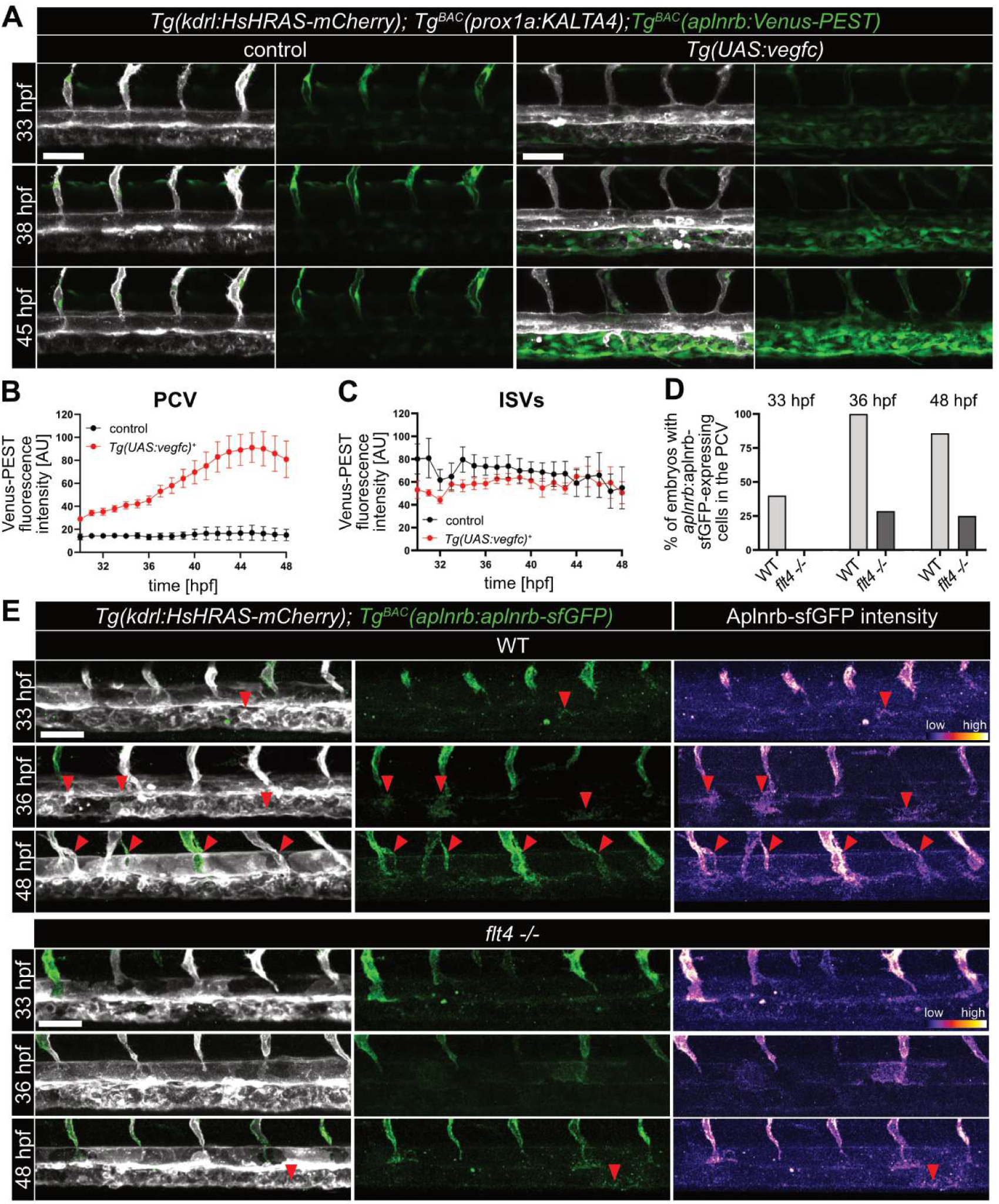
Overexpression of Vegfc induces *aplnrb* expression specifically in the PCV. **A** Still images of timelapse movies of *Tg(kdrl:HsHRAS-mCherry); Tg^BAC^(aplnrb:Venus-PEST); Tg(prox1a:KALTA4); Tg(UAS:vegfc)* embryos from 30 – 48 hpf. The broad overexpression of Vegfc leads to enhanced expression of *aplnrb:*Venus-PEST restricted to the PCV. **B, C** Quantification of the corrected total cell fluorescence of *aplnrb:*Venus-PEST in the PCV and in four ISVs (average values) corresponding to panel A. Overexpression of Vegfc leads to enhanced *aplnrb:*Venus-PEST expression accelerating at around 37 hpf when LEC emerge from the PCV while the expression in the ISVs remains constant. Data are represented as mean ± SEM. **D** Quantification of *aplnrb:*Aplnrb-sfGFP expression in the PCV of wild type and *flt4* mutant embryos between 33 to 48 hpf corresponding to panel E. In wild type siblings, Aplnrb-sfGFP-expressing cells begin to appear at 33 hpf and show robust expression by 36 hpf. In contrast, *flt4* mutants rarely exhibit Aplnrb-sfGFP-expressing cells at 36 hpf or later. n(WT)= 10 (33 hpf), 5 (36 hpf) and 7 (48 hpf); n(*flt4 -/-*)= 6 (33 hpf), 7 (36 hpf) and 8 (48 hpf). **E** Confocal images of *Tg(kdrl:HsHRAS-mCherry)*; *Tg^BAC^(aplnrb:aplnrb-TagRFP-sfGFP)* wild type and *flt4* mutant embryos at 33, 36 and 48 hpf. Aplnrb-sfGFP fluorescence intensity is visualized with a fire LUT in the right panel. Scale bars: 50 µm. See also Figure S6 and Video S2.

To investigate the mechanism through which Vegfc signaling regulates *aplnrb* expression, we examined the role of ERK signaling, a known downstream effector of Vegfc–Flt4 signaling during lymphangiogenesis [7, 40–42]. We tested whether direct ERK activation could induce *aplnrb*:Venus-PEST expression by mosaic expression of *UAS*:Map2k2b-p2A-tdTomato [7], a constitutively active form of MEK, in *Tg(fli1a:Gal4FF); Tg^BAC^(aplnrb:Venus-PEST)* embryos (Figure 7A, B). ECs expressing *UAS:*Map2k2b-p2A-tdTomato exhibited a threefold increase in *aplnrb:*Venus-PEST expression compared to controls (Figure 7C).

**Figure 7.**
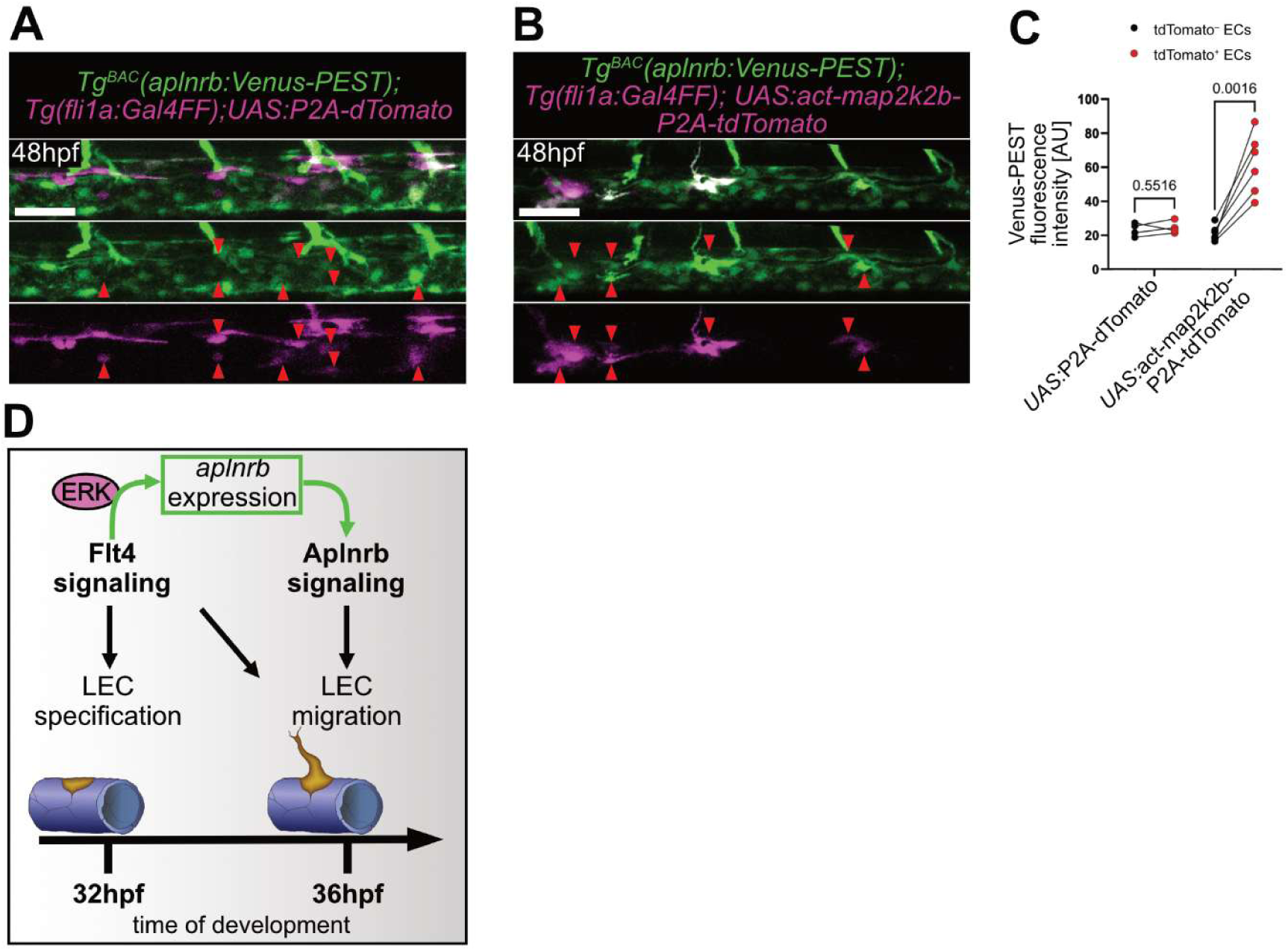
ECs with constitutively active ERK signaling show enhanced *aplnrb* expression in the PCV. **A** Confocal images of *Tg(fli1a:Gal4FF); Tg^BAC^(aplnrb:Venus-PEST)* larvae injected with *UAS:P2A-dTomato* plasmid and **B** *UAS:act-map2k2b-P2A-tdTomato* at the one-cell stage at 48 hpf. ECs of the PCV expressing *dTomato* (**A**, red arrowheads) show on average no changes in *aplnrb* reporter expression. ECs of the PCV expressing *map2k2b* (tdTomato-positive ECs, **B,** red arrowheads), the activated kinase of ERK, show higher *aplnrb* reporter expression. **C** Paired quantification of fluorescence intensity (gray values) of ECs in the PCV that are tdTomato-negative (black data points) and tdTomato-positive (red data points). Each paired value represents the measurement in one embryo. ECs expressing *UAS:*Act-Map2k2b-P2A-tdTomato show a three-fold increase in *aplnrb:*Venus-PEST fluorescence. n(*UAS:P2A-dTomato*)= 4, n(*UAS:act-map2k2b-P2A-tdTomato*)= 6. *p*-values were calculated by paired t-test. **D** Model for Vegfc–Flt4 and Apelin-Aplnrb signaling pathway interaction during lymphangiogenesis. Vegfc–Flt4 signaling is required for LEC specification (yellow cell) and regulates *aplnrb* expression via ERK signaling. Aplnrb signaling is required for LEC sprouting from the PCV. Scale bars: 50 µm.

Together, our findings support a model in which ERK modulates *aplnrb* expression, coordinating endothelial migration during lymphangiogenesis. This highlights that Apelin and Vegfc signaling coordinate and establishes Apelin signaling as a key effector of Vegfc–Flt4 signaling during lymphangiogenesis (Figure 7D).

## Discussion

Our results show that Apelin signaling plays an indispensable role during lymphangiogenesis in the zebrafish trunk by promoting the migration of LEC precursors. In *apln* mutants, lymphatic vessels are entirely absent in the trunk, in contrast to the rather mild phenotype during arterial sprouting [21, 22], indicating that LECs are particularly sensitive to the loss of Apelin signaling. Nevertheless, *apln* mutant zebrafish as well as Apln knockout mice are viable and fertile [43–45]. Interestingly, when challenged in a model of secondary lymphedema, Apln knockout mice showed reduced lymphangiogenesis in the skin. Consistently, dermolipectomy samples from patients with secondary lymphedema showed decreased APLN expression [19], supporting a conserved and context-dependent role for Apelin in lymphangiogenesis across species. In zebrafish, overexpression of Apelin leads to ectopic sprouting of ECs from the PCV, further reinforcing Apelin’s pro-migratory function. Among the two known Apelin ligands in zebrafish, mutants for the ligand *apela* showed no phenotype during lymphangiogenesis, whereas *apln* mutants exhibit the most severe phenotype. On the receptor level, *aplnra* mutants appeared phenotypically normal, while *aplnrb* mutants exhibit only mild lymphatic defects. This discrepancy between ligand and receptor mutants could be due to transcriptional adaptation [46] between the paralogous *aplnra* and *aplnrb*, which may act redundantly under loss-of-function conditions. Conclusively, *aplnra; aplnrb* double mutants completely lacked lymphatic sprouts and phenocopied the *apln* mutant phenotype. Therefore, while our data highlights Aplnrb as the dominant receptor in this context, a role for Aplnra during lymphangiogenesis cannot be entirely ruled out. Notably, our *aplnrb* reporter analysis revealed temporally restricted Aplnrb expression in a subset of sprouting ECs, particularly within the venous niche of the PCV known to give rise to LECs [33]. These findings position Apelin signaling as a key regulator of LEC migration and a promising target for modulating lymphatic vessel formation.

While Vegfc–Flt4 signaling in zebrafish is essential for both LEC specification and sprouting [3, 7, 12, 47], our data show that Apelin signaling is dispensable for LEC fate determination but is indispensable for their migration out of the PCV. Importantly, our data further indicate that Apelin signaling primarily regulates the initial migratory behavior of LECs emerging from the PCV. In *apln* mutants, LECs are correctly specified but accumulate within the PCV, resulting in an increased number of LECs in the PCV despite the absence of lymphatic sprouts. Interestingly, both Apelin and Vegfc are expressed within arterial ISVs and by fibroblasts at the horizontal myoseptum, a key site for guiding LEC migration [22, 32]. In addition, Vegfc processing components are expressed in a tightly controlled spatiotemporally pattern, which allows localized Vegfc maturation and signaling [32, 47]. Vegfc from the DA is required for the budding of lymphatic progenitors from the PCV, whereas Vegfc from fibroblasts at the horizontal myoseptum is required for the subsequent directional migration of LECs [32]. Similarly, Apelin is also expressed as a pro-peptide that can be cleaved into different isoforms, with the short Apelin-13 exhibiting the highest bioactive potency [48]. Therefore, one might hypothesize that a comparably fine-tuned spatiotemporal control of *apln* expression and processing is necessary during lymphangiogenesis to guide LECs. This might also explain why re-expression of *apln* solely in the vasculature of *apln* mutants did not result in a complete rescue. Interestingly, expression of *apln* in neural progenitors resulted in very sparse lymphatic vessel formation in *apln* mutants. It has been shown that motor neurons in zebrafish direct lymphatic progenitors to the horizontal myoseptum [49]. Similarly, in mice peripheral nerves expressing CXCL12 guide dermal lymphatic vessels via CXCR4 [16]. In zebrafish, Cxcl12a is expressed at the horizontal myoseptum and Cxcr4 by migrating LECs [50], contributing to patterning but not initial sprouting of LECs [50]. Thus, neural progenitor-derived Apelin may help refine or stabilize directional cues needed for LEC migration. Despite overlapping expression domains, re-expression of Apelin in *vegfc:*Gal4-expressing cells failed to rescue lymphatic defects in *apln* mutants. This may be due to insufficient strength of the *vegfc* promoter, incorrect timing, or the requirement for Apelin expression specifically in ISV ECs to initiate sprouting. This highlights the spatial and temporally sensitive requirement for Apelin signaling during lymphangiogenesis.

Apelin is known as a pro-migratory signaling cue in ECs [51, 52]. Strikingly, ubiquitous overexpression of Apelin led to excessive formation of ectopic endothelial extensions from the PCV. Of interest, combined haploinsufficiency of *flt4* and *apln* resulted in a complete failure of lymphatic vessel formation, suggesting a synergistic role for both pathways. Despite this strong functional interaction between Apelin and Vegfc–Flt4 signaling, we did not detect transcriptional regulation of *vegfc* or *flt4* by Apelin signaling. Importantly, we found that Vegfc overexpression and ERK overactivation both led to *aplnrb* upregulation, linking growth factor signaling to GPCR responsiveness. Previous studies [19, 23], have proposed that Apelin–Aplnr signaling functions in parallel to the Vegfc–Vegfr3/Flt4 pathway during lymphangiogenesis. While our findings do not contradict this model, they reveal an additional regulatory layer in which Vegfc–Vegfr3/Flt4 signaling acts upstream of Apelin receptor expression in LECs, establishing Apelin responsiveness during the sprouting phase. Of note, our model does not exclude the existence of additional pathways downstream of Vegfc that contribute to lymphatic sprouting. Vegfc–Flt4 signaling is known to activate both ERK and AKT pathways [7, 8]. However, pharmacological inhibition of AKT has been shown to have no effect on lympho-venous sprouting from the PCV, and expression of a constitutively active AKT transgene does not rescue venous sprouting in *flt4* mutants [7]. Consistently, we found that ERK activation is sufficient to induce *aplnrb* expression, further supporting a specific role for the ERK pathway in establishing Apelin responsiveness. Importantly, our findings do not exclude a downstream role for AKT signaling following Apelin receptor activation. Previous work by Kim et al. investigated AKT signaling downstream of Apelin signaling during lymphangiogenesis [23], whereas our experiments specifically address the upstream regulation of *aplnrb* expression. Thus, our data place ERK signaling upstream of *aplnrb* induction during the establishment of Apelin responsiveness, while AKT may still function downstream of activated Apelin signaling to regulate endothelial behavior. We therefore propose a model in which Vegfc signaling primes LECs via ERK-mediated induction of *aplnrb*, enabling these cells to respond to Apelin’s pro-migratory cues.

Lymphangiogenesis is a critical process for fluid homeostasis, immune cell trafficking, and tissue repair [53]. Dysregulated lymphatic development is implicated in pathological conditions, including lymphedema and cancer metastasis [54]. Our results reveal that Apelin signaling plays a central role during lymphangiogenesis. Importantly, Apelin signaling specifically regulates LEC migration, offering a potential therapeutic target that avoids interfering with the broader roles of Vegfc signaling. Notably, APLN-VEGFC mRNA delivery has been proposed in a preclinical trail as a therapeutic option for secondary lymphedema and is planned to enter Phase I/II clinical trail [19]. Taken together, our findings provide novel insights of Apelin signaling during lymphangiogenesis. Further work may uncover new strategies for treating lymphatic diseases through targeted modulation of LEC migration by modulating Apelin signaling.

## Material & Methods

### Zebrafish husbandry

All zebrafish housing and husbandry were performed under standard conditions in accordance with institutional (University of Marburg (UMR)) and national ethical and animal welfare guidelines approved by the ethics committee for animal experiments at the Regierungspräsidium Gießen, Germany, as well as the Federation of European Laboratory Animal Science Associations (FELASA) guidelines [55]. Embryos were raised in E3 medium, staged by hpf at 28.5°C [56] and for image acquisition, the medium was supplemented with 0.1% (w/v) propylthiouracil (PTU) from 24 hpf. See table below for lines used (Table 1).

**Table 1.** Zebrafish resource table.

| Zebrafish line | Source | Identifier |
| --- | --- | --- |
| *Tg(fli1:EGFP)<sup>y1</sup>* | Lawson & Weinstein, 2002 [57] | ZFIN: y1Tg |
| *Tg(fli1a:nEGFP)<sup>y7</sup>* | Roman et al., 2002 [58] | ZFIN: y7Tg |
| *Tg(kdrl:EGFP)<sup>s843</sup>* | Beis et al., 2005 [59] | ZFIN: s843Tg |
| *Tg(kdrl:HsHRAS-mCherry)<sup>s896</sup>* | Chi et al., 2008 [60] | ZFIN: s896Tg |
| *Tg(kdrl:mVenus-gmnn)<sup>ncv3</sup>* | Fukuhara et al., 2014 [30] | ZFIN: ncv3Tg |
| *Tg(-5.2lyve1b:DsRed)<sup>nz101</sup>* | Okuda et al., 2012 [61] | ZFIN: nz101Tg |
| *Tg<sup>BAC</sup>(apln:Venus-PEST)<sup>mr15</sup>* | Malchow et al., 2024 [21] | ZFIN: mr15Tg |
| *Tg<sup>BAC</sup>(aplnrb:Venus-PEST)<sup>mr13</sup>* | Qi et al., 2022 [62] | ZFIN: mr13Tg |
| *Tg<sup>BAC</sup>(aplnrb:aplnrb-TagRFP-sfGFP)<sup>bns309</sup>* | Helker et al., 2020 [22] | ZFIN: bns309Tg |
| *Tg<sup>BAC</sup>(prox1a:KALTA4,4xUAS-E1B:TagRFP)<sup>nim5</sup>* | van Impel et al., 2014 [31] | ZFIN: nim5Tg |
| *Tg(fli1a:Gal4FF)<sup>ubs4</sup>* | Zygmunt et al., 2011 [63] | ZFIN: ubs4Tg |
| *Tg(-2.1zic2b:Gal4-VP16)<sup>mr29</sup>* | Malchow et al., 2024 [21] |  |
| *Tg<sup>2BAC</sup>(vegfc:Gal4FF)<sup>bns270</sup>* | Parab et al., 2021 [64] | ZFIN: bns270Tg |
| *Tg(5xUAS:EGFP)<sup>nkuasgfp1a</sup>* | Asakawa et al., 2008 [65] | ZFIN: nkuasgfp1aTg |
| *Tg(UAS-E1B:NTR-mCherry)<sup>c264</sup>* | Davison et al., 2007 [66] | ZFIN: c264Tg |
| *Tg(UAS:LIFEACT-GFP)<sup>mu271</sup>* | Helker et al., 2013 [67] | ZFIN: mu271Tg |
| *Tg(UAS:apln)<sup>mr24</sup>* | Malchow et al., 2024 [21] |  |
| *Tg(UAS:vegfc)<sup>mr32</sup>* | This manuscript |  |
| *Tg(hsp70l:apln)<sup>mu269</sup>* | Helker et al., 2020 [22] | ZFIN: mu269Tg |
| *apela<sup>br13</sup>* | Chng et al., 2013 [26] | ZFIN: zf439 |
| *apln<sup>mu267</sup>* | Helker et al., 2015 [68] | ZFIN: mu267 |
| *aplnra<sup>mu296</sup>* | Helker et al., 2015 [68] | ZFIN: mu296 |
| *aplnrb<sup>mu281</sup>* | Helker et al., 2015 [68] | ZFIN: mu281 |
| *flt4<sup>hu4602</sup>* | Hogan et al., 2009 [11] | ZFIN: hu4602 |

### Generation of transgenic zebrafish lines

The *Tg(UAS:vegfc)^mr32^* line was generated by co-injection of 30 pg of plasmid DNA containing 10xUAS-zfvegfc-Tol2pACryGFP [3] and 30 pg Tol2 mRNA in wild type (AB) embryos at the one-cell stage [69]. Embryos expressing GFP in the lens were raised to adulthood and screened for germline transmission to generate a stable transgenic line.

### Whole mount *in situ* hybridization

*In situ* hybridizations were performed as previously described [67, 70]. The following probes were synthesized as published: *vegfc* [47] *, flt4* [71]*, etsrp* [67] and *aplnrb* [68]. Images were acquired using a Nikon SMZ18 binocular and DS-Fi3 camera.

### Induction of ubiquitous Apelin overexpression

Overexpression of Apelin by using *Tg(hsp70l:apln)^mu269^* was performed as previously described [21]. If not otherwise indicated, *Tg(hsp70l:apln)* embryos were subjected to a heat shock for 60 min at 37°C at 24 hpf. After image acquisition, larvae were genotyped for the *mu269Tg* allele. Wild type siblings served as control.

### Rescue experiments

For rescue experiments of *apln* mutants the following matings were performed: 1.) *apln^mu267/wt^; Tg(hsp70l:apln)* fish were mated to *apln^mu267/wt^* and the offspring was subjected to a heat shock at 24 hpf (global overexpression). *apln^mu267/wt^; Tg(UAS:apln)* fish were either mated to 2.) *apln^mu267/wt^; Tg^2BAC^(vegfc:Gal4FF); Tg(fli1a:EGFP)* (fibroblast and DA overexpression), or 3.) *apln^mu267/wt^; Tg(-2.1zic2b:Gal4-VP16); Tg(UAS-E1B:NTR-mCherry); Tg(fli1a:EGFP)* (neural overexpression), or 4.) *apln^mu267/wt^; Tg(fli1a:Gal4FF); Tg(fli1a:EGFP)* (vascular overexpression). Offspring was imaged at 120 hpf at a confocal microscope followed by genotyping. *apln^mu267/wt^; Tg(UAS:vegfc)* fish were crossed to *apln^mu267/wt^; Tg^BAC^(prox1a:KALTA4,4xUAS-E1B:TagRFP); Tg(5xUAS:EGFP)*. Offspring was imaged at 120 hpf at a confocal microscope followed by genotyping.

### Genotyping of zebrafish larvae

Embryos were genotyped by high-resolution melt (HRM) analysis [72] and analyzed using Eco ProStudy 5.2 (PCR max) or by PCR. Genotyping method and primers used are summarized below (Table 2).

**Table 2.**
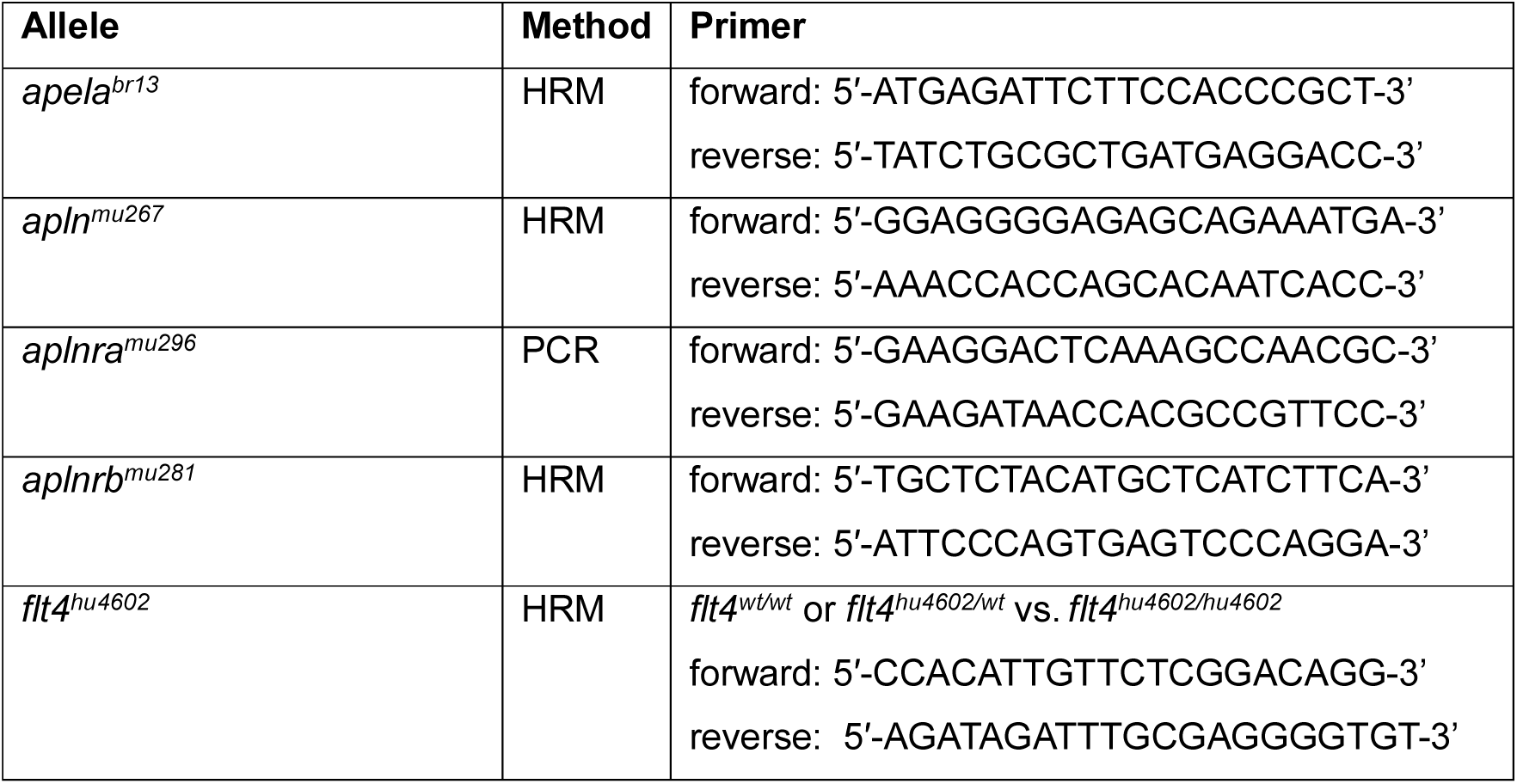
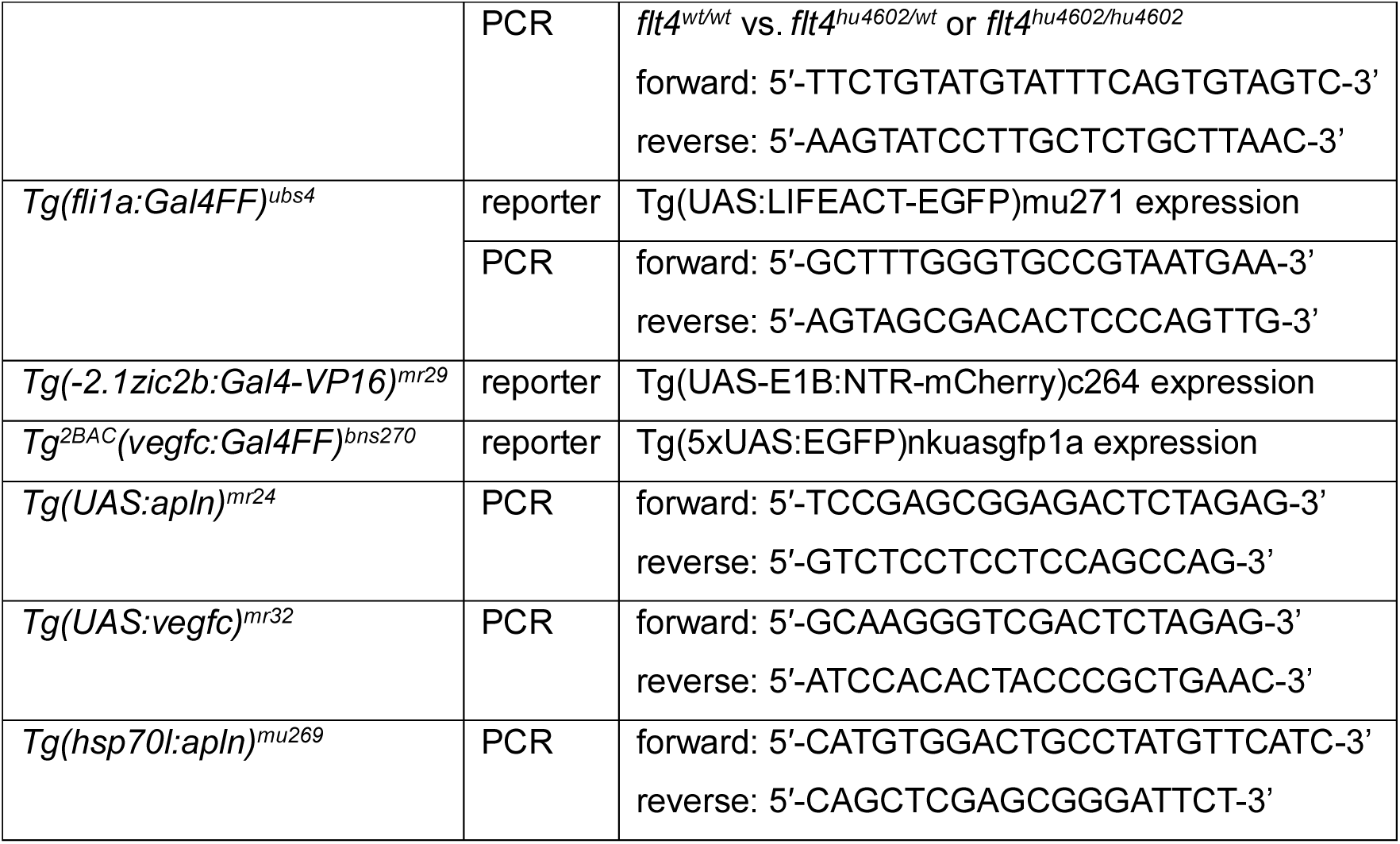
List of primers for genotyping of zebrafish.

### Mosaic ERK activation

For mosaic activation of ERK in ECs, we injected either a control plasmid containing *UAS:P2A-dTomato* or a plasmid containing *UAS:act-map2k2b-P2A-tdTomato* flanked by Tol2 sites along the Tol2 mRNA into *Tg(fli1a:Gal4FF); Tg(UAS:LIFEACT-GFP); Tg^BAC^(aplnrb:Venus-PEST)* embryos at the one-cell stage. The *map2k2b* contains point mutations resulting in a constitutively active form of MEK and therefore over-activates ERK signaling [7, 73].

### Live Imaging

Zebrafish embryos were mounted in 0.5% low melt agarose and for timelapse imaging in 0.3% normal agarose. E3 and agarose were supplemented with tricaine (19.2 mg/liter). Fluorescence images were acquired with a Nikon SMZ18 binocular and DS-Fi3 camera. Confocal images were acquired with a Leica Stellaris 8 confocal microscope equipped with HC FLUOTAR L VISIR 25x/0.95 WATER objective. For timelapse imaging, an Oko-lab incubator was set to 28.5°C. Images were analyzed and using Imaris 9.7.2 (Oxford Instruments) or ImageJ2 2.14.0 (NIH).

### *apln* mutant phenotype analysis

For vISV sprout quantification from widefield images, vISV and aISV were counted across somites 6 to 20 in the trunk region. Secondary sprouts across somites 10 to 15 were counted from confocal images. Venous sprouts were identified based on their connection to aISVs [13] and only those sprouts originating from the PCV and extending beyond the dorsal site of the DA were included in the calculations.

To categorize the TD formation, somites 10 to 15 were analyzed and the percentage of segments containing a TD was calculated. Phenotypes were categorized as follows: 100%: normal TD; <0% to <100%: partial TD; 0%: no TD.

EC numbers and distribution in the PCV and DA were quantified using Bitplane Imaris 9.7.2 software. EC nuclei were manually counted in 5 somites for both vessels.

Vessel diameter and somite width were measured in five somites per embryo by drawing a perpendicular line and using the measurement tool in ImageJ.

For quantification of venous and lymphatic sprouts emerging from the PCV, venous and lymphatic sprouts were distinguished based on their anatomical position and behavior during sprouting by using the *Tg(kdrl:HsHRAS-mCherry) line*. Venous sprouts were defined as sprouts that established a connection with an arterial intersegmental vessel, thereby contributing to the formation of venous intersegmental vessels. In contrast, lymphatic sprouts were defined as sprouts that migrated dorsally alongside the ISVs or reached the horizontal myoseptum without forming a venous anastomosis.

mVenus-gmnn-positive ECs were quantified by counting Venus expressing cells within the PCV in 5 somites.

### Fluorescence intensity measurements

Venus-PEST fluorescence intensity was quantified from confocal z-stacks using Fiji/ImageJ. For each experiment, embryos were imaged with identical acquisition settings, including laser power, detector gain, pinhole, z-step size, and exposure settings, to allow direct comparison between samples. Maximum intensity projections were generated from comparable z-ranges covering the analyzed vascular region. To quantify the Venus-PEST fluorescence intensity, the corrected total cell fluorescence (CTCF) across the entire PCV or the average of four ISV were calculated as previously described [74]. In embryos injected with *UAS:act-map2k2b-P2A-tdTomato*, regions of interest (ROIs) were defined using the polygon selection tool in ImageJ by outlining each tdTomato-expressing cell within the PCV as well as the full PCV area. To isolate fluorescence from tdTomato-positive cells, the gray values of the Venus fluorescence of all corresponding ROIs were measured using the OR (Combine) function. For quantification of tdTomato-negative background fluorescence within the PCV, the XOR function was applied to exclude tdTomato-positive ROIs from the PCV measurement.

### Statistics and reproducibility

All experiments were performed using at least three independent biological replicates (independent clutches). Comparable numbers of embryos were analyzed per experimental condition. Imaging and phenotypic analyses were performed prior to genotyping, and genotypes were determined afterward in a blinded manner. Statistical analysis was performed in Prism 10.3.1 (GraphPad). Respective tests and n numbers are indicated in the figure legends. We identified outliers using the Robust regression and Outlier removal (ROUT) method with a threshold of 1%. We used two-tailed unpaired Student’s t test to compare between two means and One-way analysis of variance (ANOVA) with Dunnett’s multiple comparison test for comparison of multiple conditions. For non-normally distributed data, One-way ANOVA with Kruskal-Wallis test was used for comparison of multiple conditions. Categorical data were compared using the Fisher’s exact test. If not indicated otherwise, values are shown as mean ± SD. *p*-values of <0.05 were regarded as significant.

## Resource Availability

Requests for further information and resources should be directed to and will be fulfilled by the lead contact, Christian S.M. Helker.

All unique/stable reagents generated in this study are available from the lead contact with a completed materials transfer agreement.

## Statements and Declarations

### Ethics approval

All animal experiments were performed in accordance with institutional (University of Marburg (UMR)) and national ethical and animal welfare guidelines approved by the ethics committee for animal experiments at the Regierungspräsidium Gießen, Germany, as well as the Federation of European Laboratory Animal Science Associations (FELASA) guidelines.

### Author contributions

Conceptualization, C.S.M.H. and F.L.; Methodology, C.S.M.H. and F.L.; Formal Analysis, F.L., J.E. and E.B.; Investigation, F.L., E.B., E.E., J.M.; Resources, C.S.M.H.; Writing – Original Draft, C.S.M.H. and F.L., Writing – Review & Editing, C.S.M.H., F.L., J.E., L.W.W; Visualization, F.L., J.E.; Supervision, C.S.M.H.; Funding Acquisition, C.S.M.H.

## Acknowledgements

We would like to thank Sabine Fischer, Regina Löchel and Peter Braun for technical support and Stefan Baumeister for the schematic model. Microscopy was performed with the support of the Centre for Advanced Light Microscopy (CALM), Philipps-University Marburg funded by the German Research Foundation (DFG, project number: 446988475). Image analysis was performed with the support of the Core Facility “Zelluläre Bildgebung“, Philipps-University Marburg. This work was supported by the German Research Foundation (DFG, project number: 506858860).

## Supplemental Information

**Video S1.** Timelapse movie of *Tg(kdrl:HsHRAS-mCherry); Tg(fli1a:nEGFP)* larvae from 30 hpf to 46 hpf. In *apln* mutant larvae, venous endothelial cells (ECs) within the posterior cardinal vein (PCV) fail to initiate sprouting compared to sibling controls. Dark blue spheres mark ECs remaining within the PCV. Light blue spheres indicate ECs contributing to venous sprouts, and magenta spheres indicate ECs contributing to lymphatic sprouts.

**Video S2, related to Figure 6**. Timelapse movie of *Tg(kdrl:HsHRAS-mCherry); Tg^BAC^(aplnrb:Venus-PEST); Tg(prox1a:KALTA4); Tg(UAS:vegfc)* embryos from 30 – 48 hpf. Vegfc overexpression induces *aplnrb*:Venus-PEST expression in the posterior cardinal vein (PCV).

## Supplemental Figures

**Figure S1.**
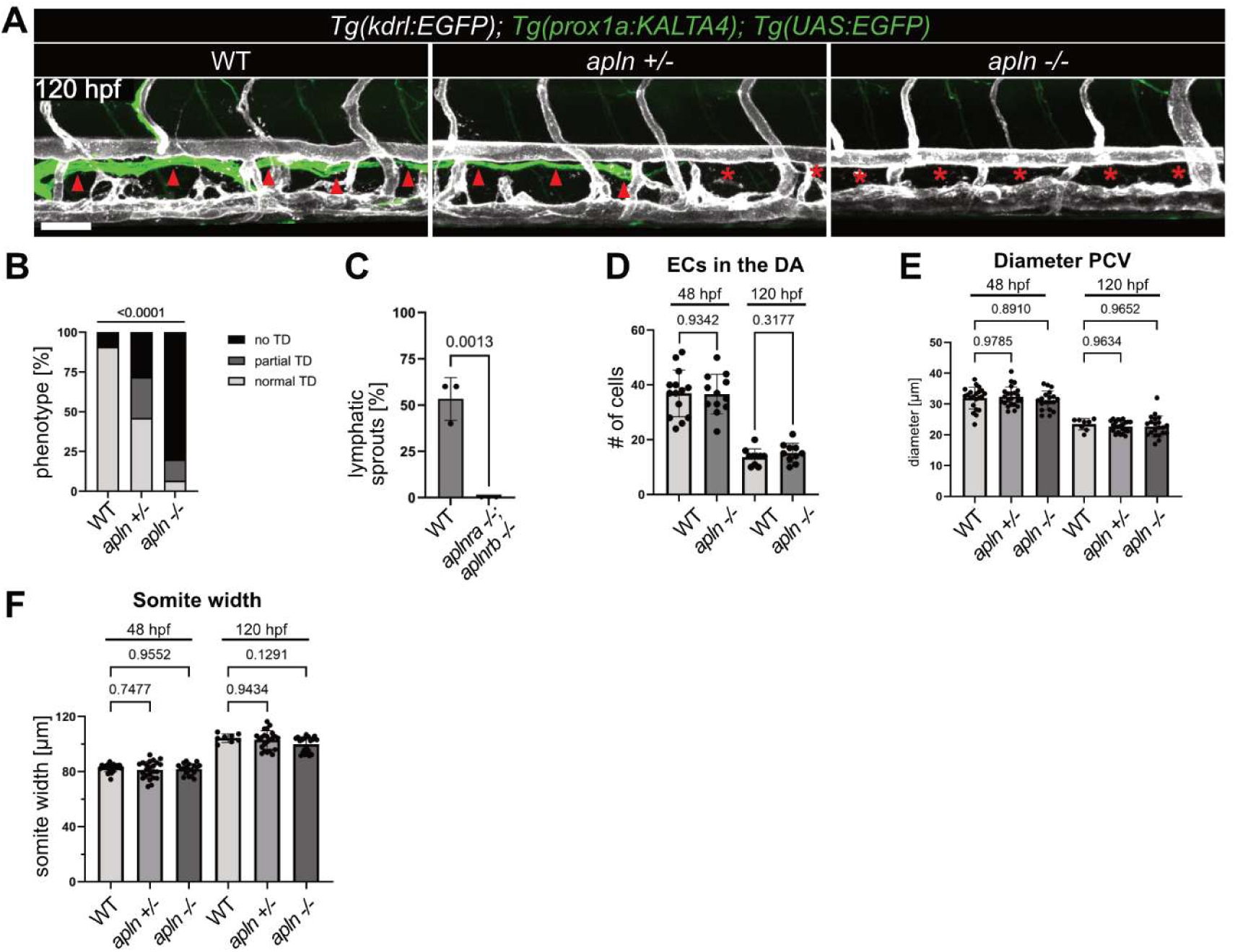
related to Figure 1 – Apelin signaling is required for lymphatic development and *apln* mutants have an increased cell number in the PCV. A Fluorescence images of *Tg(kdrl:EGFP)*; *Tg(prox1a:KALTA4); Tg(UAS:EGFP)* larvae at 120 hpf. *apln (+/-)* mutants exhibit partial TD formation (red arrowheads and red asterisks) and *apln (-/-)* mutants do not exhibit a TD (red asterisks). **B** Quantification of the observed TD phenotype. n(WT)= 11; n(*apln +/-*)= 39; n(*apln -/-*)= 15. **C** Quantification of lymphatic sprouts of *Tg(fli1a:EGFP)* larvae at 48 hpf. n(WT)= 3; n(*aplnra -/-; aplnrb -/-*) = 3. *p*-value was calculated by Student’s t-test. **D** Quantification of number of ECs in the DA at 48 and 120 hpf. There is no significant difference between *apln* mutants and WT siblings. **E** Quantification of the PCV diameter at 48 and 120 hpf. There is no significant difference between *apln* mutants and WT siblings. **F** Quantification of average somite width at 48 and 120 hpf. There is no significant difference between *apln* mutants and WT siblings. E and F: 48 hpf: n(WT)= 22, n(*apln+/-*)= 23, n(*apln -/-*)= 19. 120hpf: n(WT)= 8, n(*apln +/-*)= 23, n(*apln -/-*)= 21. Scale bars: 50 µm.

**Figure S2.**
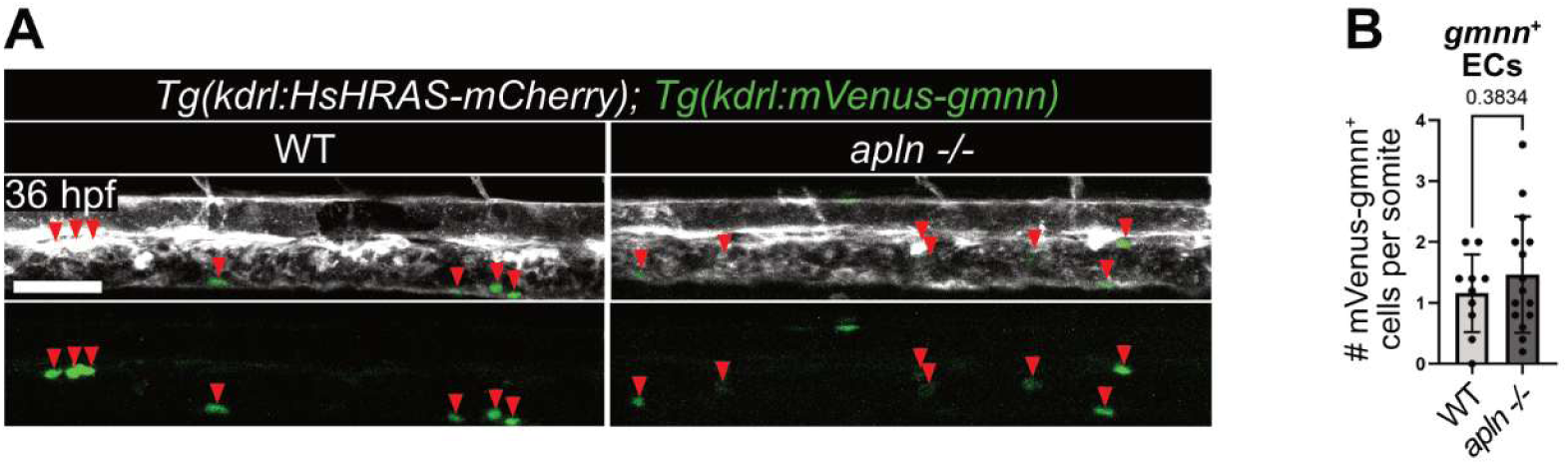
related to Figure 1 – *apln* mutants do not show increase mitotic activity in the PCV. **A** Confocal images of *Tg(kdrl:HsHRAS-mCherry); Tg(kdrl:mVenus-gmnn)* embryos at 36 hpf. Red arrowheads indicate *kdrl:*mVenus-gmnn-expressing cells in the PCV. **B** Quantification of *kdrl:*mVenus-gmnn-expressing cells (mitotic cells) in the PCV per somite at 36 hpf. Compared to their siblings, there is no significant difference in *apln* mutants. n(WT)= 10, n(*apln -/-*)= 15. Data are represented as mean ± SEM. *p*-values were calculated by Student’s t-test. Scale bar: 50 µm.

**Figure S3.**
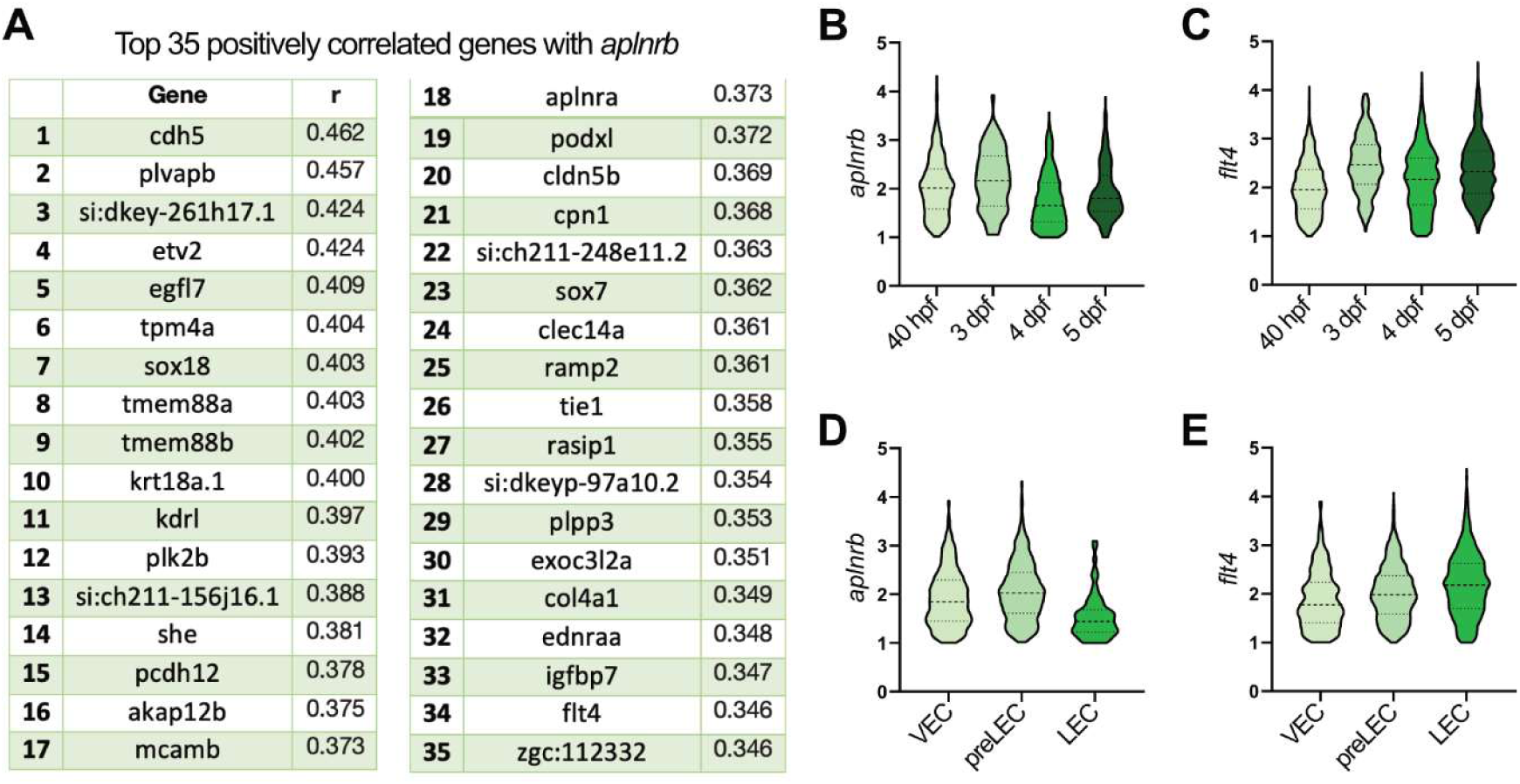
related to Figure 2 – Single cell data reveal *aplnrb* expression in early lymphatic cells. **A** List of top 35 *aplnrb* positively correlated genes in the vasculature [34]. Among these are the lymphatic progenitor marker *etv2*, the pro-migratory genes including egfl7 and pro-lymphatic genes such as *sox18*, *tie1* and *flt4*. **B-E** Analysis of gene expression from the single cell data of lymphangiogenesis [4]. **B** *aplnrb* is predominantly expressed during early lymphangiogenesis and **C** in preLECs while **D** *flt4* is expressed throughout lymphatic development and **E** strongest in LECs.

**Figure S4.**
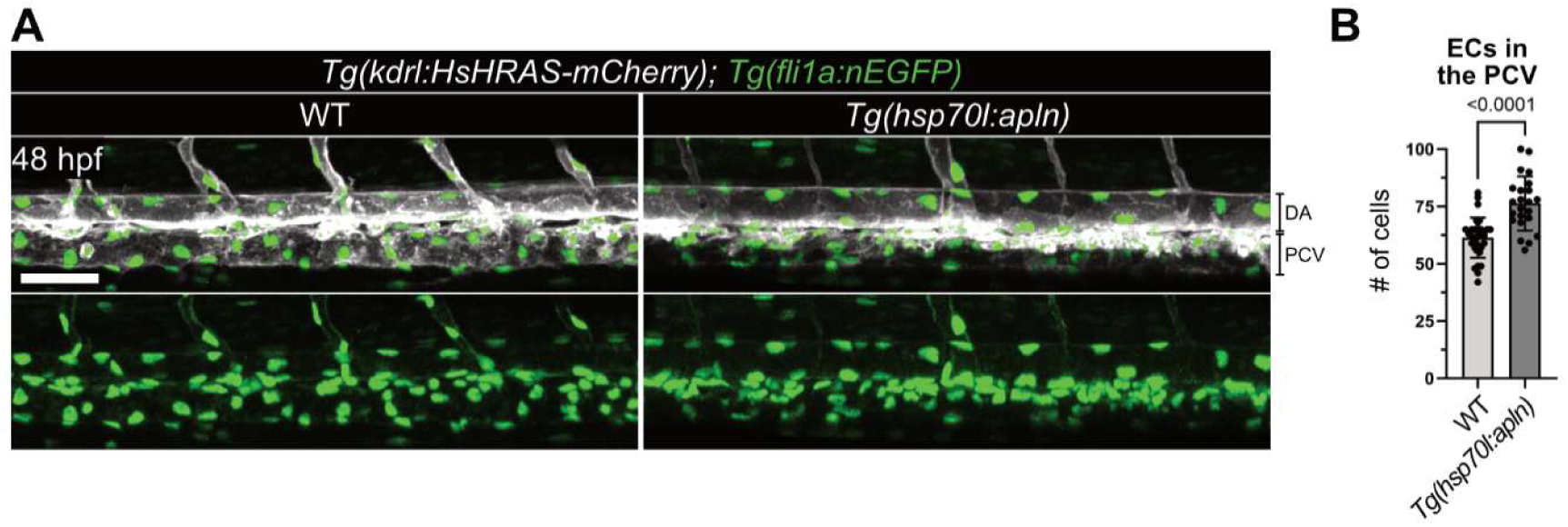
related to Figure 3 – Overexpression of Apelin leads to an increased cell number in the PCV. **A** Quantification of global Apelin overexpression in *Tg(hsp70l:apln)* embryos at 48hpf. Apelin overexpression by *Tg(hsp70l:apln)* leads to a higher cell number in the PCV. **B** Quantification of cell number. *p*-value was calculated by Student’s t-test. n(WT)= 32, n*Tg(hsp70l:apln)*= 23.

**Figure S5.**
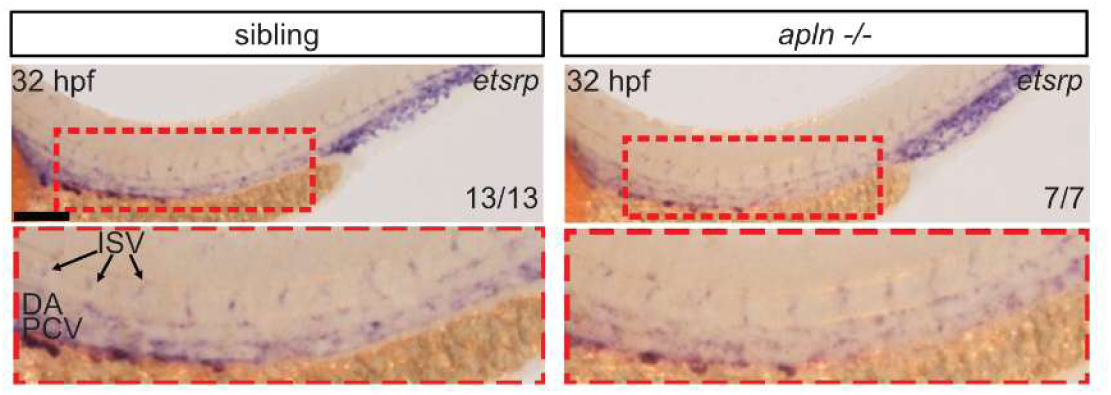
related to Figure 5 – Apelin does not regulate *etsrp* mRNA expression. Whole mount *in situ* hybridization against *etsrp* of *apln* mutants and siblings at 32 hpf. Magnification of red boxed area below. Compared to siblings, *apln* mutants exhibit a similar *etsrp* expression pattern. Scale bar: 100 µm. ISV: intersegmental vessel.

**Figure S6.**
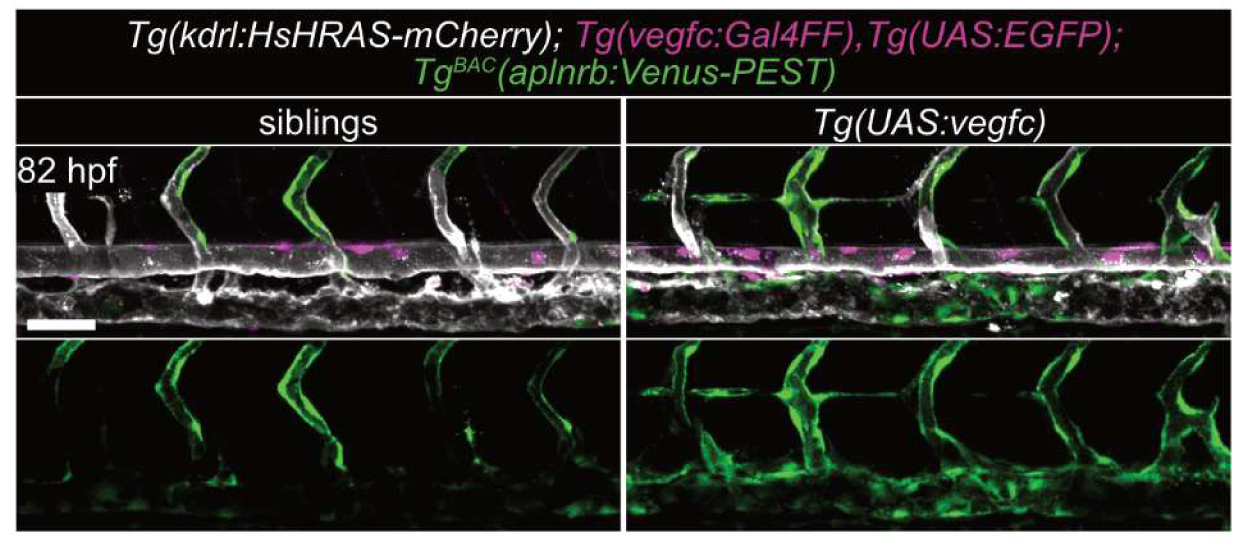
related to Figure 6 – Overexpression of Vegfc induces *aplnrb* expression specifically in the PCV. Confocal images of *Tg(kdrl:HsHRAS-mCherry); Tg(vegfc:Gal4FF); Tg(UAS:Vegfc); Tg^BAC^(aplnrb:Venus-PEST)* larvae at 82 hpf. Overexpression of Vegfc in Vegfc-expressing cells is sufficient to enhance expression of *aplnrb:*Venus-PEST.

